# Experimental evolution toward extinction in a molecular host-parasite system

**DOI:** 10.1101/2025.08.31.673314

**Authors:** Kohtoh Yukawa, Tomoaki Yoshiyama, Ryo Mizuuchi, Norikazu Ichihashi

## Abstract

Theoretical studies have proposed that coevolution with parasitic replicators plays a critical role in the evolution of primitive life; however, experimental verification of the potential outcomes of such coevolutionary dynamics remains limited. We previously conducted a co-evolutionary experiment using an RNA-protein replication system that resulted in the spontaneous diversification of host and parasitic RNAs into five distinct lineages with robust co-replication. Here, we report contrasting evolutionary outcomes from a second long-term co-evolutionary experiment. Using a droplet flow reactor system with increased dilution frequency over 5000 h (1,600 generations), we observed reduced diversity and frequent extinctions in later experimental stages. Co-replication assays of RNA clones revealed that the primary cause of this diversity loss was the shortened reaction time resulting from frequent dilution. Further analysis of RNA clones that emerged during evolution suggested that the frequent extinctions resulted from the appearance of highly competitive parasite species and the dominance of host species that exhibited reduced replication ability. These findings demonstrate that co-evolution between host and parasitic replicators can result in diversity loss and frequent extinctions depending on dilution conditions, highlighting the critical role of environmental parameters, such as dilution ratio and frequency, in enabling primitive replicators to evolve sustainably toward the emergence of life.

## Introduction

Many scenarios for the origins of life propose that self-replicating molecules, such as self-replicating RNA, acquired the ability of Darwinian evolution on the ancient Earth and subsequently evolved through increasing complexity and diversity to approach extant life (Szathmáry and John Maynard 1995; Joyce 2002; Koonin and Martin 2005; Yarus 2011; Pross and Pascal 2013; Higgs and Lehman 2015; Goldenfeld et al. 2017; Adamski et al. 2020; Brunk and Marshall 2021). However, whether and how self-replicating molecules can spontaneously develop diversity and complexity through evolutionary processes remains largely unknown.

One promising approach involves experimental evolution using replicating molecules capable of evolution. The first experimental molecular evolution was reported in 1967 by Spiegelman’s group, in which phage genomic RNA was replicated by purified phage RNA replication enzyme (replicase) across multiple generations (Mills et al. 1967). Subsequently, other RNA/DNA replication systems were developed and employed for experimental evolution (Breaker and Joyce 1994; Wright and Joyce 1997). Although these systems demonstrated biologically relevant phenomena, including niche differentiation (Voytek and Joyce 2009) and adaptation to inhibitory chemicals (Kramer et al. 1974; Paegel and Joyce 2010), the typical evolutionary outcome was dominance by the fastest replicators under given conditions, with no observed diversification.

Contrasting results were obtained when a translation system was integrated with Spiegelman’s RNA replication system (designated the translation-coupled RNA replication system) in our previous study (Ichihashi et al. 2013). In this system, RNA encoding an RNA replicase is replicated by the replicase translated from itself in a reconstituted E. coli translation system (Fig. 1a). A notable feature of this system is the emergence of parasitic RNAs that have lost the replicase-encoding sequence while retaining the replicase recognition sequence through RNA recombination. These parasitic RNAs replicate by exploiting “host” RNAs that retain the replicase gene but replicate significantly faster than host RNAs due to their reduced size. To prevent excessive amplification of parasitic RNAs, the system was compartmentalized within water-in-oil droplets. Through serial dilution cycles, the RNA sustained replication and continued evolving. In a previous long-term evolutionary experiment more than 500 generations, we observed co-evolution of host and parasitic RNAs through an evolutionary arms race (Furubayashi et al. 2020), wherein the ancestral host RNA diversified into at least five distinct lineages—three host lineages and two parasite lineages—forming an interdependent replication network (Mizuuchi et al. 2022).

**Figure 1:**
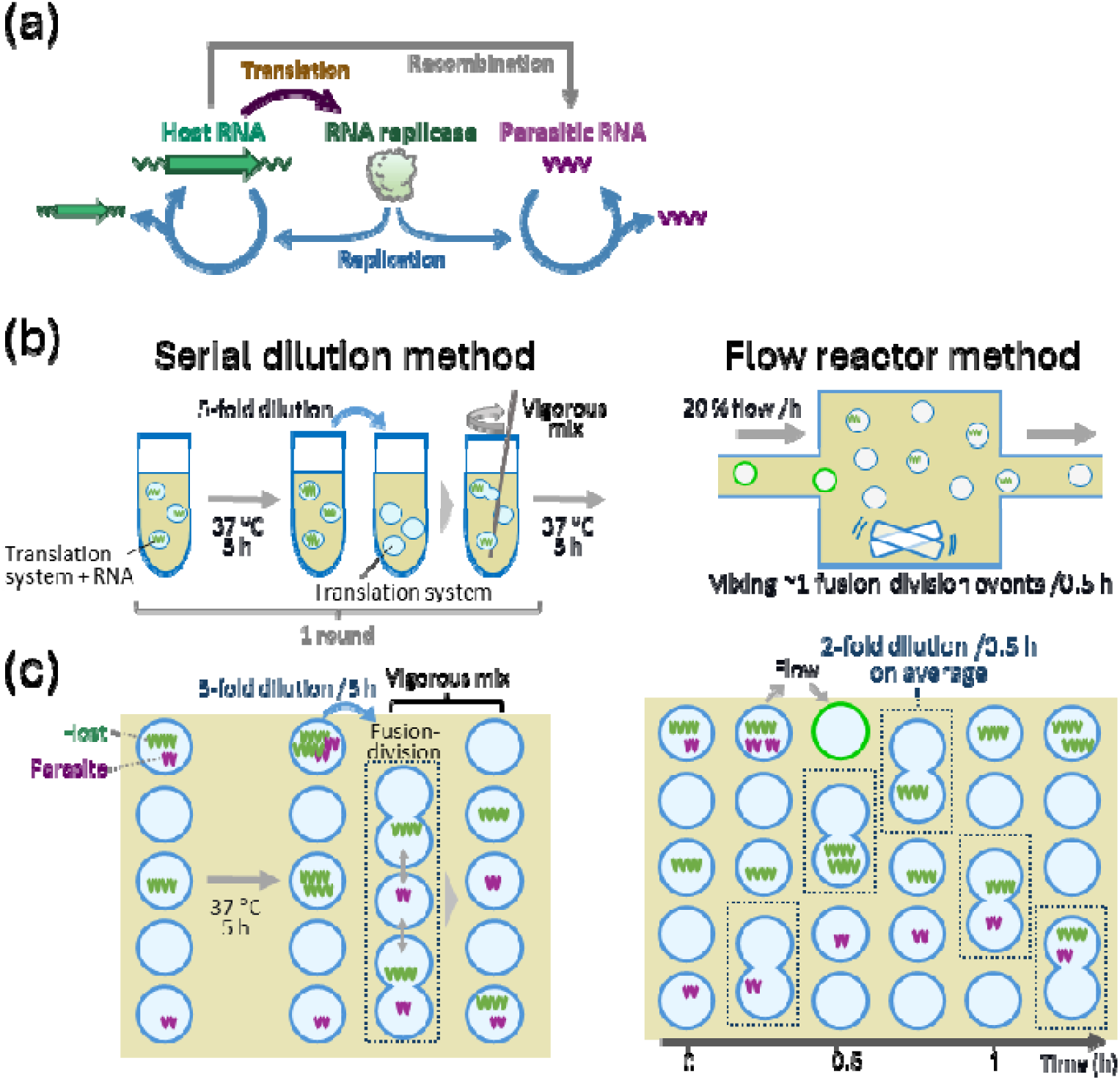
A long-term experiment using a flow reactor. (a) Scheme of translation-coupled RNA replication. RNA replicase is translated from a host RNA and replicates both host and parasitic RNAs. Parasitic RNA occasionally appears from a host RNA through recombination. Both host and parasitic RNAs accumulate mutations by replication error and evolve according to the Darwinian principle. (b) Differences between the previous serial dilution (SD) and the new flow reactor (FR) methods. In the SD method, water-in-oil droplets containing the host and parasite RNA and the translation system were incubated for 5 h at 37°C for RNA replication. Then, an aliquot of the droplets was diluted 5-fold with new droplets containing the translation system, vigorously mixed to prepare new droplets with completely mixed contents, and incubated for 5 h at 37°C again for the next round of replication. In the FR method, water-in-oil droplets were incubated at 37°C and stirred for 3 s every 60 s, which allows approximately one fusion-division event/0.5 h, corresponding to 2-fold dilution of droplet contents /0.5 h. New droplets containing the translation system were supplied constantly at a rate of exchange of 20% of the total volume per hour. (c) Schematic image of droplets in each condition. RNAs in the droplets experienced 5-fold dilution /5 h in the SD method, while 2-fold dilution /0.5 h on average in the FR method. These two methods can be compared based on dilution ratio and frequency.

One factor contributing to the distinct evolutionary outcomes in the translation-coupled RNA replication system may be the emergence of parasitic entities. Parasitic molecules have been considered a critical obstacle for primitive self-replicators to overcome before the emergence of life, as parasites inevitably arise in self-replicating systems (Iranzo et al. 2016) and disrupt interdependent or cooperative replication networks (Eigen and Schuster 1978; Maynard Smith 1979). Several solutions to this problem have been proposed, including compartmentalization, whether permanent or transient (Bresch et al. 1980; Szathmáry and Demeter 1987; Bianconi et al. 2013; Matsumura et al. 2016; Furubayashi and Ichihashi 2018; Laurent et al. 2019; Blokhuis et al. 2020), limited dispersion (Boerlijst and Hogeweg 1991; Boerlijst and Hogeweg 1995; Cronhjort and Blomberg 1997; McCaskill et al. 2001; Szabó et al. 2002; Sardanyés and Solé 2007; Shay et al. 2015), molecular kin selection (Levin and West 2017), and physical linkage evolution (Levin et al. 2020). Even when the problem of parasite amplification is resolved through these mechanisms, how hosts and parasites co-evolve remains an open question.

Theoretical studies have predicted diverse coevolutionary outcomes (reviewed in (Buckingham and Ashby 2022)) and some are related to primitive replicators: increased host diversity and complexity (Takeuchi and Hogeweg 2008; Zaman et al. 2014; Hickinbotham et al. 2021; Seoane and Solé 2023), enhanced host variation with stabilized oscillation dynamics (Decaestecker et al. 2013), or extinctions of a host species due to reduced population sizes during oscillations (Gokhale et al. 2013). While many coevolutionary experiments using extant organisms have been conducted (reviewed in (Brockhurst and Koskella 2013)), these studies employed relatively short time scales compared to the theoretical frameworks for primitive replicators mentioned above, and theoretical predictions regarding diversification and extinction dynamics remain insufficiently verified. In our aforementioned long-term coevolutionary experiment using the translation-coupled RNA replication system, we always observed stable coexistence and divergence of host and parasitic RNAs in three evolutionary experiments (Kamiura et al. 2022; Mizuuchi et al. 2022). However, whether this evolutionary outcome consistently occurs in this system or whether alternative outcomes are possible under different conditions remains unknown.

In this study, we conducted another longer-term evolutionary experiment up to 1,600 generations under increased serial dilution frequency using a flow reactor system and observed a contrasting coevolutionary outcome: evolution toward reduced diversity and periodic extinction. During the early stage of evolution, host and parasitic RNA continued replicating with oscillatory dynamics similar to our previous experiment; however, in later stages, host RNA concentrations frequently declined substantially and became undetectable (i.e., underwent extinction). We also observed that the diversified host and parasite lineages present in the initial population were eliminated shortly after starting this evolutionary experiment, with only one host and one parasite lineage persisting in later stages. Short-term replication assays using cloned host and parasitic RNAs revealed that loss of diversity resulted from frequent dilution, and periodic extinctions were caused by the appearance of highly competitive parasites and the reduction in host replication ability.

## Result

### Evolutionary experiment with a flow-reactor system

For another evolutionary experiment utilizing an automated format, we employed a previously developed droplet flow reactor system (Yoshiyama et al. 2018). In this system, droplets containing the translation system were supplied to the reactor at a constant rate. Within the reactor, droplets containing host and parasitic RNAs were stirred with a magnetic stirrer at a specified frequency to induce fusion and division events, thereby mixing the contents and enabling RNA to continue replication. Details of the flow reactor system are presented in Fig. S1.

The difference between the new flow reactor (FR) condition and the previous manual serial dilution (SD) condition is illustrated in Figs. 1b and 1c. Under the SD condition, droplets were incubated for 5 h at 37°C to induce RNA replication inside, and then a portion of the droplets was diluted 5-fold with new oil phase containing droplets that included the translation system. The diluted droplets were vigorously mixed using a homogenizer to achieve nearly complete mixing of the droplet contents, then incubated for the subsequent replication cycle. Under this condition, both the droplets themselves and their contents were diluted simultaneously 5-fold every 5 h.

Under the FR condition, the droplets in the reaction tank were continuously diluted with new droplets that included the translation system to exchange 20% of the tank volume per hour, while a magnetic stirrer periodically induced droplet fusion-division at a rate of approximately one fusion-division event per 0.5 h (Yoshiyama et al. 2018). Under the FR condition, droplets and internal RNAs were diluted independently; droplets were continuously diluted through the droplet flow to exchange 20% of tank volume per hour (i.e., 1.25-fold/h), while the RNAs in a droplet were discontinuously diluted at 2-fold/0.5 h through fusion-division events with another droplet, most of which contained neither host nor parasitic RNAs according to the observed concentrations in the evolutionary experiment shown below (∼10^−1^ nM, corresponding to ∼0.01 molecule/droplet). Note that the latter discontinuous RNA dilution at a droplet fusion (2-fold/0.5 h) directly affects internal RNA concentrations and their replications, and thus we compared it with that under the SD condition in the following experiments.

Using this FR system, we conducted a long-term evolutionary experiment employing the RNA population from round 155 of our previous evolutionary experiment under the SD condition, in which RNA had already diversified into three host and one parasite lineage. Host RNA concentrations were quantified by reverse transcription followed by quantitative PCR (RT-qPCR). Figure 2 presents host RNA concentrations for both the previous evolutionary experiment under the SD condition (light green lines) and the current experiment under the FR condition (dark green lines). Under both conditions, host RNA concentrations exhibited oscillatory dynamics; however, the average RNA concentration under the FR condition was lower than under the SD condition. Host RNA continued to be detected for approximately 2000 h under the FR condition, but was undetected (i.e., extinct) after 2264 h (top panel in Fig. 2). We restarted the experiment using preserved droplets from 1920 h (indicated by the red dotted line). Host RNA replication then persisted slightly longer, suggesting that extinction was stochastic, but again became undetectable after 2600 h (second panel from top). In this way, we continued the evolutionary experiment until 5024 h and experienced extinctions eight times in total. Extinctions occurred frequently in later experimental stages, particularly around 2000, 3000, and 4500 h, suggesting that RNA populations evolved during these periods may possess characteristics that induce them to extinction.

**Figure 2:**
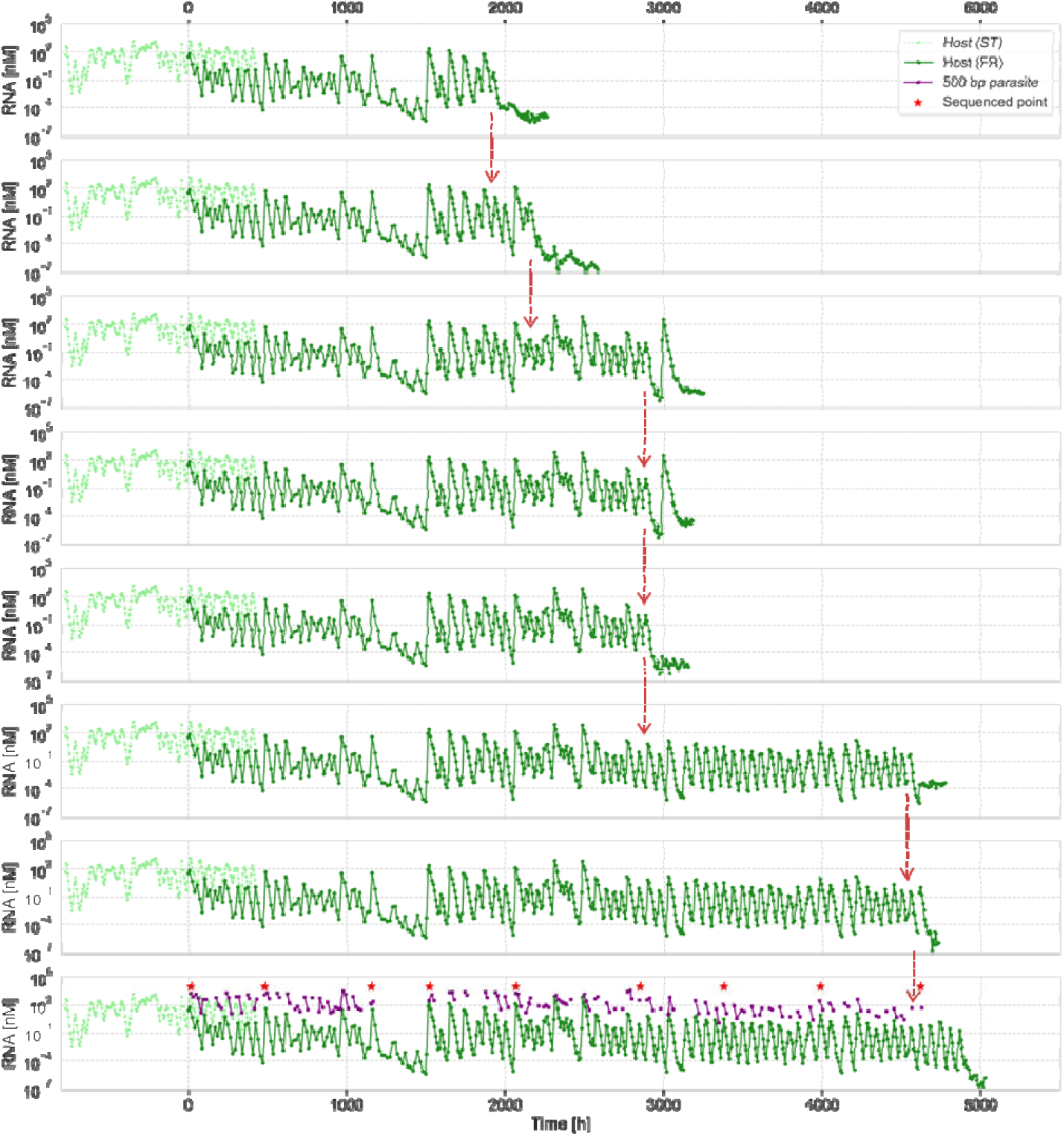
Trajectories of host RNA concentration during the FR evolutionary experiment until each extinction. The host RNA concentrations were measured by RT-qPCR using host-specific but lineage-nonspecific primers. The results of the previous evolutionary experiment under the SD condition from rounds 0 to 240 are shown in light green, assuming that one round was equivalent to five hours. The new evolutionary experiment under the FR condition was started from the RNA population of round 155 under the SD condition, and the results are shown in deep green. When the host dropped below the detection limit multiple times and did not recover, the experiment was restarted from the stocked sample at an earlier time point (indicated by the red dotted arrows). Parasitic RNA concentrations, measured by quantification from SDS-PAGE images, are shown in the bottom panel (purple markers). The time points used for sequence analysis are indicated by red stars.

The complete trajectory of RNA concentration for the experiment with the longest duration up to 5024 h (corresponding to approximately 1,600 generations; see Methods for calculation) is presented in the bottom panel of Fig. 2. For this experiment, we also quantified parasitic RNA concentrations (purple marks) based on band intensity following polyacrylamide gel electrophoresis (Fig. S2). The detection limit for the band intensity method is substantially higher than that of RT-qPCR employed for host RNA quantification; consequently, parasitic RNA was detectable only at elevated concentrations. Parasites were detected intermittently around host RNA concentration peaks, indicating that parasite RNA concentrations also oscillated in accordance with host oscillations. In subsequent analyses, we examined RNA populations from this longest-duration evolutionary experiment.

### Sequence analysis

To elucidate the evolutionary dynamics of hosts and parasites during the FR evolutionary experiment, we analyzed host and parasite RNA sequences at nine time points (red stars in Fig. 2) using long-read deep sequencing (PacBio Sequel) after reverse-transcription. To eliminate the effect of sequence error and quasispecies, we identified dominant mutations occurring at frequencies of 10% or higher at least one analyzed time point and classified sequences into genotypes based on the presence or absence of these dominant mutations (Figs. S3 and S4). We constructed a phylogenetic tree incorporating the three most abundant genotypes at each analyzed time point (solid lines in Fig. 3a), along with the data before round 155 of the previous evolutionary experiment (dotted lines). We also presented the frequency of each genotype as a heatmap aligned with the phylogenetic tree.

**Figure 3:**
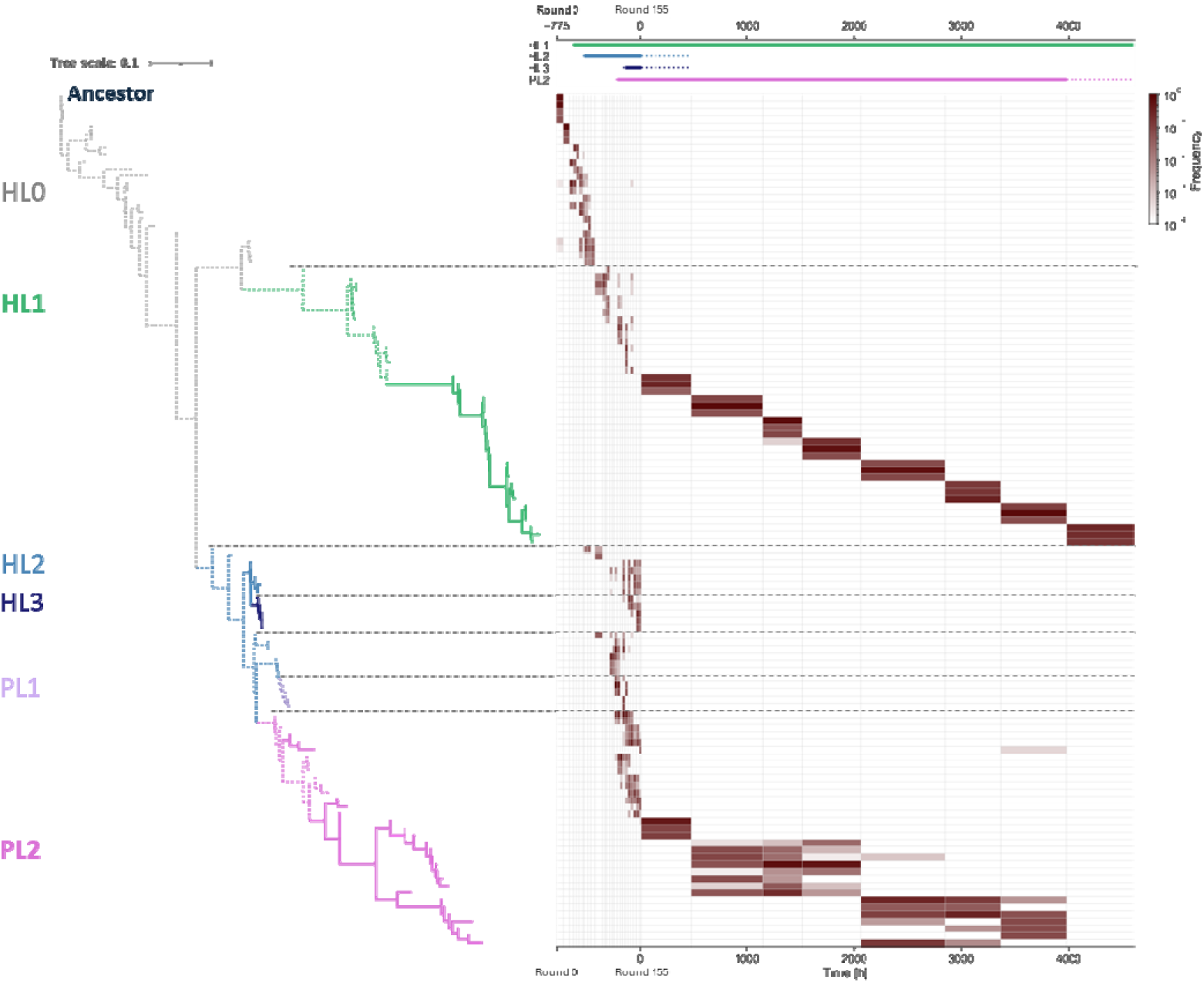
Phylogenetic tree and frequencies in the population. A phylogenetic tree was constructed using the three most frequent genotypes at each sequencing time point. The branches for the previous SD experiment (until round 155) and the new FR experiment are shown as dashed and solid lines, respectively. Different types of RNA are depicted in different colors based on the classification in a previous study (Mizuuchi et al. 2022). The heatmap shows the frequencies of the genotypes in each branch. Schematic representation of existing RNA lineages is shown on the top. Uncertain regions due to the lack of sequence data were shown in dotted lines.

At the starting point (0 h) of the FR experiment (i.e., round 155 of the SD experiment), three host lineages (HL1, HL2, and HL3) and one parasite lineage (PL2) were present (Fig. 3). However, after 480 h, HL2 and HL3 disappeared, with only one host (HL1) and one parasite (PL2) remaining detectable, indicating that diversified lineages were not sustained under the FR condition. The existence of each lineage is schematically shown at the top of the heatmap. The heatmap additionally reveals a relatively homogeneous population consisting of closely related branches until 4616 h, differing by only 2-3 mutations in the host population shown in Fig. 3. This result indicates that new distinct lineages did not emerge under the FR condition. These results under the FR condition represent a stark contrast to the previous SD experiment (Fig. S5), in which five distinct lineages emerged.

### Conditions for maintaining diversity

We investigated the differences between the FR and SD conditions that influence diversity maintenance (i.e., the number of coexisting RNA species). We chose two factors that primarily differ between the FR and SD conditions: (1) the dilution frequency, which determines the continuous reaction time that host and parasitic RNAs experiences before a dilution event, and (2) the dilution ratio at a dilution event. The dilution frequency and the dilution ratio under the SD condition correspond to once /5 h and 5-fold, respectively, because RNAs were diluted 5-fold every 5 h (Fig. 1b, left). Under the FR condition, the dilution frequency and the dilution ratio of RNAs in droplets correspond to once /0.5 h and 2-fold, respectively, because a droplet experiences one fusion-division events per 0.5 hour on average (Fig. 1b, right). A more detailed rationale for this calculation is described in the Methods section.

We hypothesized that these differences in the two factors affect the number of coexisting RNA species. To test this hypothesis, we conducted short-term replication experiments employing the serial dilution method using multiple RNA species at varying dilution frequencies and ratios, including those corresponding to the FR and SD conditions, then evaluated the number of RNA species present in the final population.

As RNA species for this experiment, we initially selected four RNA species from round 155 (HL1, 2, 3, and PL2); however, these did not exhibit stable co-replication (Fig. S6), likely because these RNAs had not yet established stable interdependencies. We therefore employed five RNA clones from a later round—three host RNAs (HL1, HL2, and HL3) and two parasitic RNAs (PL2 and PL3)—identified at round 228 in the previous study (Mizuuchi et al. 2022), in which four of these five RNA species (excluding HL3) demonstrated co-replication under the SD condition (dilution frequency of once /5 h and dilution ratio of 5-fold).

Before conducting the experiments, we investigated the effect of dilution frequency and ratio on the number of co-replicable RNA species using mathematical modeling and simulation. This mathematical model was based on a previous study (Kamiura et al. 2022). The model comprised three sequential steps: 1) RNA replication reactions dependent on the number of RNA molecules within each droplet, 2) random removal and supply of droplets, and 3) random fusion-division of droplets (see Methods for details). In these simulations, we varied the dilution frequencies for RNA replication (once/0.5 h, /1 h, and /5 h) and the dilution ratio (2, 5, 10, and 100-fold. Other parameters, including the replication rate of each RNA species, were derived from experimentally determined values (Mizuuchi et al. 2022). We executed 50 rounds of the replication-dilution-fusion-division cycle 20 times. Representative RNA concentration dynamics are presented in Fig. 4a. The average number of remaining RNA species in final populations is displayed in the upper left triangle of Fig. 4c.

**Figure 4:**
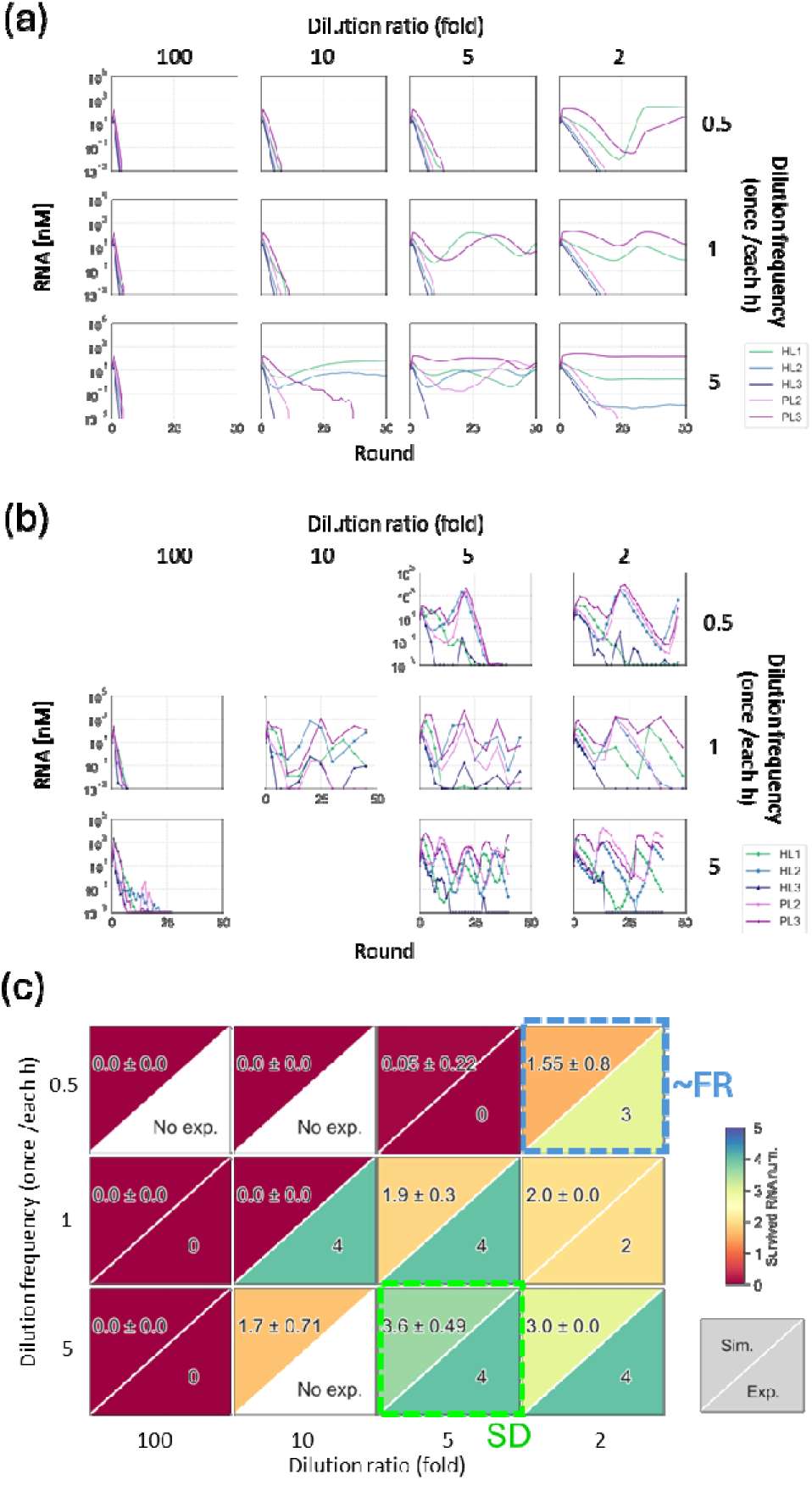
Effect of dilution rate and reaction time on the coexistence of multiple host and parasitic RNA. Short-term serial dilution experiments with five types of RNA species (three hosts and two parasites) were performed for 40 – 50 rounds at different dilution frequencies and ratios using both computer simulations (a) and experiments (b). (c) Summary of the number of RNA species in the final population in the simulation (top left, average of 20 runs with standard deviation) and the experiment (bottom right).

To validate these results, we conducted nine short-term serial dilution experiments at varying dilution frequencies and ratios and quantified the concentration of each RNA species using specific RT-qPCR (Fig. 4b). The number of maintained RNA species in the final 40–50 rounds is presented in the bottom right panel of Fig. 4c with the standard deviations. The number of remaining RNA species was significantly dependent on both dilution frequencies and ratios, with simulation and experimental results exhibiting similar trends. At frequent dilution and larger dilution ratios (upper-left region of Fig. 4c), RNA species rarely persisted, whereas at infrequent dilution and smaller dilution ratios (lower-right region), greater numbers of RNA species were maintained. Specifically, under the previous SD evolutionary condition (once/5h and 5-fold dilution, green dotted square), 3.6–4 RNA types were sustained. Under the condition corresponding to the FR evolutionary experiment (twice /h and 2-fold dilution, blue dotted square), only 1.6–3 RNA species persisted, consistent with the loss of pre-existing RNA diversity during the FR evolutionary experiment shown in Fig. 3, where two RNA lineages remained. These results suggest that reduced diversity under the FR condition can be primarily attributed to frequent dilution because another difference under the FR condition, reduced dilution ratio, should increase the diversity at the dilution frequency of twice/h based on the data of Fig. 4c.

### Cause of extinction

We investigated the underlying cause of host RNA disappearance during later stages of the evolutionary experiment. Such disappearance may be directly attributed to reduced host RNA concentrations at the bottom of the oscillation dynamics, which likely results in stochastic removal of all host RNA molecules from the reactor. If this hypothesis is correct, host RNA concentrations at the bottom should decrease in the later stage of evolution. To test this, we plotted RNA concentrations at each bottom of the oscillation in Fig. 5a (blue points), which were approximately 10□³ nM during 0–1000 h and decreased to 10□ nM during 1000–2000 h and 3000–4600 h. Beyond 4600 h, the bottom concentrations decreased further.

**Figure 5:**
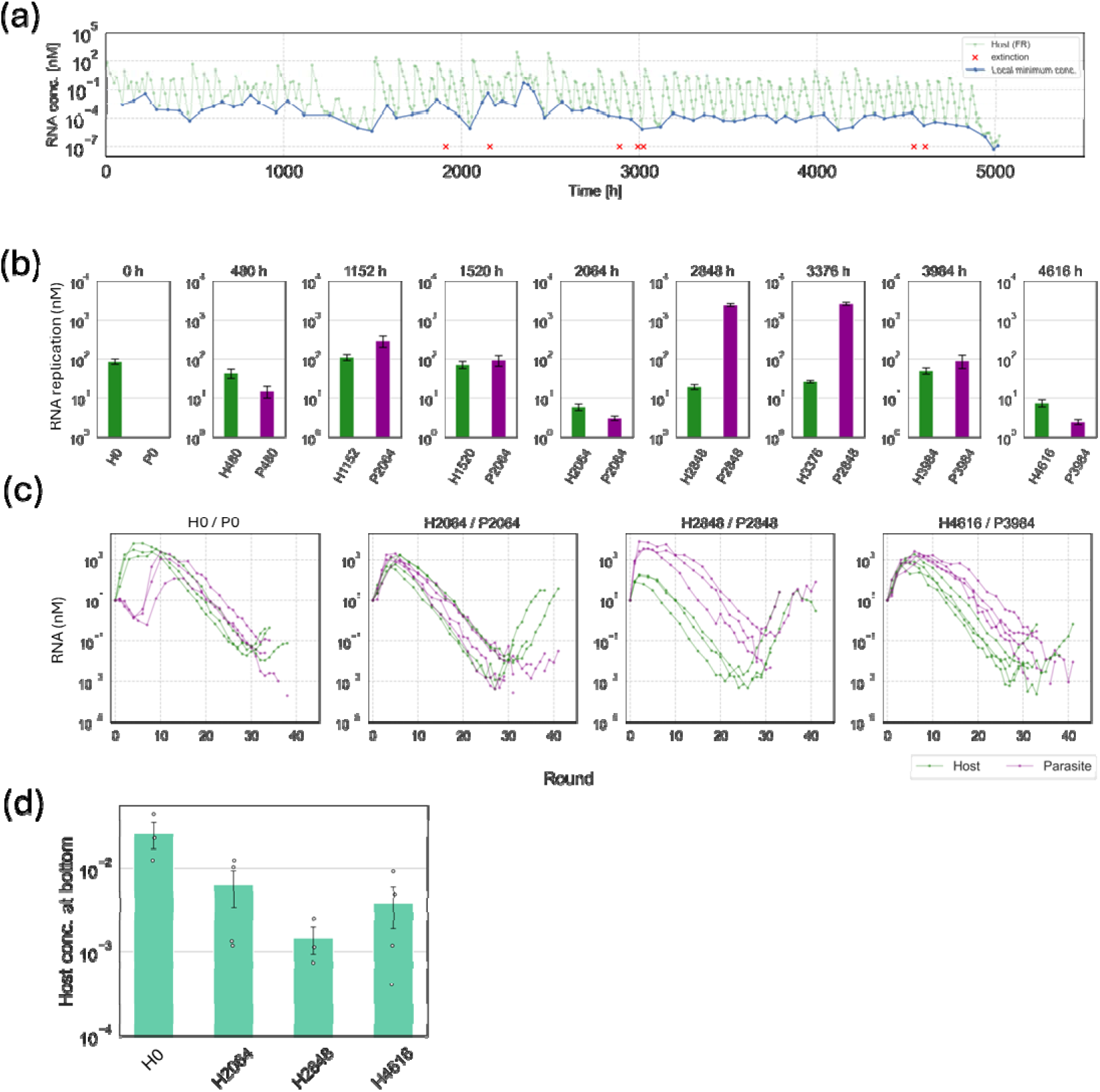
Characterization of RNA species that appeared in the FR evolutionary experiment. (a) Trajectory of the host RNA concentrations at the bottom of oscillation. (b) Single-round competition experiments with a host and parasitic RNA clones that dominated at the indicated time points. Each RNA pair (1 nM each) was incubated in the translation system at 37 °C for 1 h, and the host and parasite concentrations were measured using RT-qPCR. (c) A short-term serial dilution experiment using the host and parasite RNA pairs (10 nM each) at 0, 2026, 2848, and 4616 h under the condition close to FR condition (dilution frequency of once /0.5 h and dilution ratio of 2-fold) for 38–40 rounds. The average host RNA concentrations at the bottom are shown in (d) with standard errors (N = 3 or 4).

Next, we investigated whether evolved host and parasite RNAs in later stages exhibited characteristics that reduced host concentrations at the bottom of oscillation. We selected the most frequent host sequences at nine time points (0, 480, 1152, 1520, 2064, 2848, 3376, 3984, and 4616 h) and prepared individual RNA clones. For parasite, we selected five major sequences that were identical during 1152–2064 h and 2848–3376 h because similar sequences were sustained for relatively long periods. At 4616 h, parasitic RNA was undetectable; therefore, we employed the sequence from 3984 h. Using these host-parasite pairs from each time point (1 nM each), we conducted single-round competition experiments and quantified RNA replication (Fig. 5b).

For the initial host-parasite pair before the FR evolutionary experiment, the host replicated selectively without parasite amplification (Fig. 5b, 0 h). At 480–1520 h following initiation of the FR evolutionary experiment, parasites were amplified to levels equivalent to host replication. During 1520–2064 h, replication of both host and parasite declined. At 2848 and 3376 h, host replication increased modestly while parasite replication increased substantially. Parasite replication subsequently decreased at 3984 h, with both host self-replication activity and parasite replication declining at 4616 h. We identified two notable patterns in host-parasite replication relationships: (1) increased parasite replication (e.g., at 480 and 2848 h) and (2) reduced replication for both host and parasite (e.g., at 2064 and 4616 h).

The subsequent question was how these characteristics of evolved hosts and parasites resulted in reduced host RNA concentrations at the bottom of oscillations. In the evolutionary experiments, extinctions were frequently observed around 2000, 3000, and 4600 h. Host and parasite RNAs during these periods (2064, 2848, and 4616 h in Fig. 5b) also exhibited the aforementioned two notable patterns: reduced replication for both host and parasite (2064 and 4616 h), and increased parasite replication (2848 h). To test whether these two contrasting characteristics could result in lower host RNA concentrations at the bottom of oscillations, we conducted short-term replication experiments using host-parasite pairs from each time point (0, 2064, 2848, 3984 h) under the condition approximating FR condition (dilution frequency of once /0.5 h and dilution ratio of 2-fold), and quantified each host and parasite concentration. These experiments were performed independently in triplicate (Fig. 5c). The average host concentrations at the bottom are presented in Fig. 5d. The bottom concentration at 0 h exceeded 10□² nM but decreased to below 10□² nM for 2064, 2848, and 3984 h (Fig. 5d), indicating that all three evolved RNA pairs reduced the host concentrations at the bottom, consistent with frequent extinctions observed around these time periods.

### Replication assay using purified replicases

We further investigated how each host and parasitic RNA changed during evolution in more detail. We first purified each replicase encoded by the nine host species as a recombinant protein (Fig. S8). The total number of purified replicases was seven because the host RNAs at 1520 h and 2064 h or 2848 h and 3376 h have the same replicases. We then performed replication experiments with all combinations of the nine host and five parasitic RNAs (Fig. S9). Using these data, we analyzed the trajectory of the replicase activities (Fig. 6a), and the template activity (i.e., the activity to be replicated) of the host and parasitic RNAs (Fig. 6b) during evolution. Additionally, we analyzed the selectivity of host over parasite, the ratio of host replication to that of each corresponding parasite, for each replicase (Fig. 6c). We also measured the translation level of each replicase from the host RNA (Fig. 6d).

**Figure 6:**
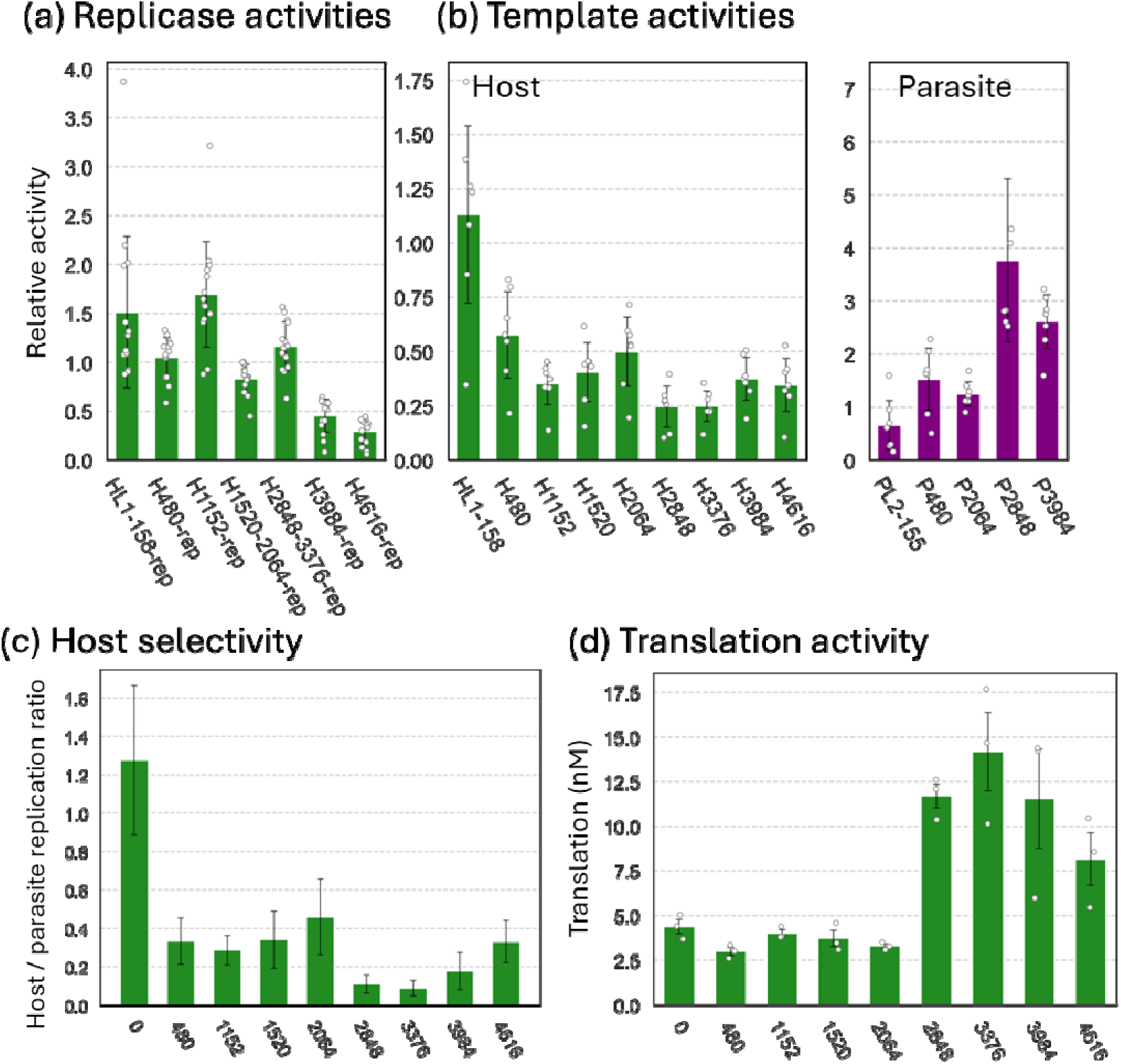
Trajectories of host and parasite activities during evolution. (a) Trajectory of the replicase activities during evolution. RNA replication results of each replicase for all host and parasite templates were averaged after normalizing template difference by dividing by the average fold replication of each template RNA. (b) Trajectory of the template activity (i.e., the activity to be replicated) of the host and parasitic RNAs during evolution. RNA replication results of each template RNA for each replicase were averaged after normalizing replicase difference by dividing by the average fold replication of each replicase. (c) Trajectory of the selectivity of the host against the parasite during evolution. The result of a host RNA replication was divided by the result of the corresponding parasite replication for each replicase. (d) Translation level of the replicase from each host RNA.

These analyses revealed at least four findings. First, both the template activity of host species and the replicase activities tended to decrease during evolution with slight variations (Fig. 6a, right and Fig. 6b), indicating that the replication ability of host species tended to decrease during evolution. Second, the template activity of the parasitic RNA, in contrast, tended to increase during evolution, indicating that more competitive parasites appeared during evolution, especially at 480 h and 2848 h. Third, the selectivity for the host tended to increase except at 480 h and 2848 h. Fourth, translation of replicase increased at 2848 h (H2848)

These changes in replication properties can explain the two aforementioned notable patterns in host-parasite replication relationships. The (1) increased parasite replication at 480 and 2848 h was probably caused by the appearance of highly competitive parasites (P480 and P2848) at these time points and by the increased translation at 2848 h. The (2) reduced replication for both host and parasite (e.g., at 4616 h) was probably caused by the combination of decreased host template activities, decreased replicase activities, and increased host specificities of the replicases.

## Discussion

Coevolution between hosts and parasites represents an important evolutionary phenomenon relevant to the origin of life and modern biology; however, experimental evidence regarding potential coevolutionary outcomes remains limited. Here, we conducted and analyzed a long-term evolutionary experiment under altered dilution frequency and demonstrated that the evolutionary outcome differed significantly from our previous evolutionary experiments, in which hosts and parasites diversified and coexisted stably in three partially independent experiments (Mizuuchi et al. 2022). Host RNA exhibited a tendency toward extinction in later evolutionary stages (Fig. 2) and lost its lineage diversity (Fig. 3). Assays using RNA clones revealed that infrequent and moderate dilution were necessary for maintaining diversity (Fig. 4). Competition assays using isolated RNA clones demonstrated that RNAs selected during later stages exhibited two contrasting properties: increased parasite replication and reduced replication for both host and parasite (Fig. 5b). Both properties were found to decrease the bottom host concentrations during oscillations (Figs. 5c and 5d), thereby increasing extinction probability. The biochemical assay using purified replicases revealed the detailed causes of these properties. The increased parasite replication was attributed to the appearance of highly competitive parasites. The reduced replication for both host and parasite was attributed to decreased host replication ability and increased host selectivity against parasites. These results indicate that host-parasite coevolution does not invariably result in robust co-replication and spontaneous diversification, as observed in our previous evolutionary experiment, but can lead to diversity loss and increased extinction susceptibility probably depending on dilution conditions. These findings underscore the critical importance of environmental parameters that influence dilution dynamics for primitive replicators to evolve sustainably toward the emergence of life.

Although evolution is generally regarded as a process that either increases or maintains the fitness of populations of organisms or molecules, it can drive population extinction in theory, a phenomenon known as evolutionary suicide, runaway evolution, or Darwinian extinction (Webb 2003; Kalle Parvinen 2005). Such evolution toward extinction has been investigated in several theoretical models (Ferrière 2000; Gyllenberg and Parvinen 2001; Parvinen 2007; Parvinen 2010; Parvinen and Dieckmann 2013), including predator-prey systems (Hiroyuki Matsuda and Peter A. Abrams 1994; Matsuda and Abrams 1994; Dieckmann et al. 1995), bacterial populations (Himeoka and Mitarai 2020), and epidemiological models (Boots and Sasaki 2003; Boldin and Kisdi 2016). However, experimental demonstration of evolution toward extinction remains limited (Rankin and López-Sepulcre 2005). In this study, we provide an experimental example of such evolution toward extinction using a molecular replication system. During the long-term evolutionary experiment under the FR condition, we observed frequent extinctions only after 2000 h (Fig. 2), suggesting that the host RNA evolved to acquire the characteristics that predispose it to extinction.

Our explanation for this counterintuitive evolutionary trajectory is as follows. First, both increase in dilution frequency and decrease in dilution ratio under the FR condition could decrease the bottom concentration according to the simulation results (Fig. S13). Furthermore, frequent dilution under the FR condition reduced the diversity of RNA species, as shown in Fig. 4. In ecological studies, reduced diversity is known to enhance population fluctuations due to the absence of buffering effects from other species (Decaestecker et al. 2013) and therefore, the loss of diversity could contribute to the reduced bottom concentration, which leads to the feasibility of extinction. However, these effects were in effect from the beginning of the evolutionary experiment, and were not sufficient to explain the frequent extinctions in the later stage of the evoultuionary experiment. The additional effects would be the appearance of less self-replicative host species and highly competitive parasite species during the evolutionary experiment (Fig. 6), both of which further reduced host RNA concentrations at the bottom of oscillations, as shown in Fig. 5c. These combinatorial effects of dilution conditions and evolution could cause stochastic extinctions in the later period.

A remaining question is why host RNA evolved to decrease its self-replication ability. Our proposed explanation is the accumulation of weakly deleterious mutations through genetic drift. This hypothesis is supported by the fact that the population size of intact host RNA became extremely small (∼10, see Methods for detailed calculation) at the bottom of oscillation dynamics. A similar phenomenon was also reported in a theoretical study, in which a reduction in population size during the host-parasite oscillation dynamics facilitated the fixation of rare mutations through genetic drift (Gokhale et al. 2013).

In summary, we observed two evolutionary phenomena during the new long-term evolutionary experiment using a droplet flow reactor system: the loss of diversity and evolution toward extinction. The loss of diversity can be explained by dilution conditions (more frequent dilution). The loss of diversity, together with frequent dilution, possibly resulted in larger amplitude in the oscillation dynamics of RNA populations, which decreased the minimum host population size. The decreased population size possibly caused the accumulation of weakly deleterious mutations in host RNA through genetic drift, which, together with the evolution of highly replicable parasites, resulted in frequent extinctions in the latter part of the evolution. As a whole, we propose that host-parasite coevolution under frequent dilution possibly results in loss of host diversity, thereby selecting for RNA species with increased susceptibility to extinction.

## Materials and Methods

### The translation-coupled RNA replication system

The translation-coupled RNA replication system used in this study originally consists of two components: a host RNA that encodes a core subunit of the Qβ RNA replicase gene with the recognition sequence, and the reconstituted translation system of *E. coli*, which contains all factors required for translation, such as ribosome, translation proteins, tRNA, NTPs, and amino acids. The components were prepared as previously described (Hagino et al. 2025). Parasitic RNAs appeared during the replication from the host RNA.

### Evolutionary experiment using a flow reactor

The droplet flow reactor system (Fig. S1) was constructed as described in a previous study (Yoshiyama et al. 2018). This system consisted of two 505 µL tanks set on magnetic stirrers. Oil phases and aqueous phases (the reconstituted translation system) were supplied by syringe pumps into the 1^st^ tank, where water-in-oil droplets were prepared by stirring with a magnetic stirrer at 2 °C. The oil phase consisted of mineral oil (Sigma) and surfactant, 2% Span 80 (Wako), and 3% Tween 80 (Wako), saturated with non-protein components of the translation system, as described previously (Mizuuchi et al. 2022). The prepared droplets were transferred to the 2nd tank, in which the droplets contained host and parasitic RNAs, and RNA replication occurred in the droplets at 37 °C. The droplets were mixed for 3 s every 60 s by stirring at 4800 rpm to induce fusion and division between the droplets. This corresponds to the low mixing condition in a previous study (Yoshiyama et al. 2018). The flow rate was 0.2/h (oil phase, 100 µL/h; aqueous phase, 1 µL/h).

At the start of the experiment, 500 µL of the oil phase and 5 µL of the translation system were added to the 1st tank, and the emulsion was prepared by stirring at 2 °C and 7200 rpm for 10 min. The second tank was filled with 505 µL of water-in-oil droplet, which was prepared by mixing droplets from round 155 of the previous SD evolutionary experiment with 4 µL of translation system and 400 µL of oil phase at 7200 rpm for 3 min using a magnetic stirrer at 2 °C. When restarting the reaction in the middle of this experiment, the water-in-oil droplets (505 µL) were added to the second tank at the restarting time point. The translation system and oil phase were replenished every 2–3 days.

### RNA measurement

To measure the host RNA concentration, the water-in-oil droplets were diluted 100-fold with 1 mM EDTA (pH 8.0) and heated at 95 °C for 5 min, followed by rapid cooling on ice. The heated sample was then used for quantitative PCR after reverse transcription (RT-qPCR) using primers 1-4 and the One-Step SYBR Prime Script PLUS RT-PCR Kit (Takara). To measure the parasitic RNA concentration, the water phase was collected from 300 µL of the water-in-oil emulsion by centrifugation at 15 krpm for 5 min. The precipitate was then washed with 120 µL of diethyl ether. After centrifugation at 10 krpm for 1 min and removal of the ether phase, the remaining aqueous phase was mixed with 50 µL of water, and RNA was purified using the RNeasy Mini Kit (QIAGEN). Purified RNA (250 ng) was applied to an 8% polyacrylamide gel, as described previously (Mizuuchi et al. 2022). As a size marker for parasitic RNA, we used a short parasite (parasite α in a previous study (Furubayashi et al. 2020)) synthesized through *in vitro* transcription. After electrophoresis at 150 V for 85 min, the gel was stained with SYBR Green II (Takara). The intensity of each band was quantified using ImageJ software.

### Sequence analysis

RNA was purified from the water-in-oil droplets as described above. Then, RNAs were reverse transcribed with PrimeScript RTase (Takara) and primers 5 and 6, followed by RNase H (Takara) treatment at 37 °C for 20 min. The cDNA was PCR-amplified using KOD FX (TOYOBO) and primers 5-6 and purified using the PureLink PCR Micro Kit (Life Technologies). The DNA corresponding to the host and parasite were then separated by gel extraction using an E-gel agarose electrophoresis system (Life Technologies), and barcode sequences were attached by PCR with KOD FX (TOYOBO) and primers 5 and 6 attached to each barcode sequence. Then, the DNA mixture was sequenced using PacBio Sequel. In PacBio Sequel, a single DNA molecule can be read multiple times owing to its circular format. We used a consensus sequence of Q > 20 to reduce sequencing error. The number of consensus sequences used was 10000 at any analyzed time point, except for parasites at 2848 h, 3376 h, and 3984 h. These three points were thought to have insufficient RNA concentrations; however, more than 2,000 sequences were obtained. Following the method described in a previous study (Mizuuchi et al. 2022), 130 dominant mutations that existed at a frequency of 10% or higher in all the analyzed samples were identified (Figs. S3 and S4). We classified the genotypes of each consensus sequence based only on the dominant mutations because each consensus sequence contains many unique mutations, probably because of the error-prone nature of RNA replication and the remaining sequencing errors. The genotypes that were observed in more than nine reads were used for the following analysis. The extracted genotypes and their frequencies are shown in the Source Data file. Phylogenetic trees were constructed using the top three most frequent genotypes from each analysis time point using the neighbor-joining method in MEGA-X version 10.2.5 (Kumar et al. 2018).

### Coexistence experiment in Fig. 4

The five RNA clones (three types of host RNA and two types of parasite RNA) found in the final round (round 228) of the previous evolutionary experiment (HL1-228, HL2-228, HL3-228, PL2-228, and PL3-228) were prepared by in vitro transcription as described previously (Mizuuchi et al. 2022). The five RNA clones were mixed at 10 nM each in the translation system, and the mixture (10 µL) and the buffer-saturated oil phase (990 µL) were vigorously mixed with a homogenizer (Polytron PT-1300D, KINEMATICA) to prepare water-in-oil droplets. The droplets were incubated at 37 °C for the indicated periods and diluted with water-in-oil droplets, which were prepared by mixing the saturated oil and translation system, at the indicated dilution ratio, using the same method without RNA addition. The diluted droplets were further incubated for the next round of replication, for the indicated periods. After repeating this serial dilution process, each RNA sample was measured using RT-qPCR with specific primers 7-16 and the One Step SYBR PrimeScript PLUS RT-PCR Kit (Takara). Although we used primers that selectively recognized only one of the five RNA clones, a low level of misrecognition occurred (usually 1/100 – 1/1000 of the legitimate recognition) (Fig. S7). This misrecognition effect was subtracted when calculating RNA concentration. For the round 155 RNA set, because sufficient accuracy could not be achieved with one-step RT-qPCR, only the plus strand was measured using primers 17-24. Primer sequences are shown in the Source Data file.

### Coexistence simulation in Fig. 4

The simulation was based on the replication-mixing-dilution model described in a previous study (Mizuuchi et al. 2022). This model consists of three steps, 1) replication, 2) dilution, and 3) fusion-division, and 300,000 droplets (compartments). 1) In the replication step, i^th^ RNA species replicate logistically in each droplet, according to the differential equations described below.

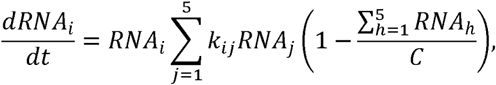

where C is a carrying capacity in a droplet (300). The replication rate constants *k_ij_*, experimentally determined previously (Mizuuchi et al. 2022), were multiplied by 0.1 in this study because the original values were too large to observe the effect of the reaction time. 2) In the next dilution step, the droplets were randomly removed according to the dilution rate, and an equal number of empty droplets were added. 3) In the subsequent fusion-division step, RNAs from two randomly selected droplets were combined and redistributed into two new droplets according to a binomial distribution to mimic the fusion-division process of droplets. This process was repeated 1.3 times the droplet number according to the previous study. Starting from an initial state in which RNAs were distributed according to a Poisson distribution with a mean of 10 molecules per droplet, the replication, dilution, and redistribution processes were repeated for 50 rounds. All parameter values used in this simulation are shown in the Source Data file.

### Rationale of the dilution frequency and rate under the SD and FR conditions

The condition corresponding to the previous evolutionary experiment (SD condition) was a dilution frequency of once /5 h and a dilution ratio of 5-fold because one round of the serial dilution cycle consisted of incubation for 5 h, followed by a 5-fold dilution.

The condition close to the new evolutionary experiment using the flow reactor (FR condition) is a dilution frequency of once /0.5 h and a dilution rate of 2-fold for the following reasons. Under the FR condition, we stirred the droplets at 4800 rpm for 3 sec/min, which corresponds to a diffusion coefficient of 0.35 /min^−1^, which means that droplets experience a fusion-division event approximately once every 0.5 h according to the data in a previous study (Yoshiyama et al. 2016) (based on Fig. 4 of the previous study). Therefore, we adopted a dilution frequency of once / 0.5h. In addition, RNA in a droplet should be diluted 2-fold in a single fusion-division event, assuming that all droplets are the same size and a fusion-division event of two droplets mixes the contents completely and produces two droplets of the same size. It should be noted that the above calculation is valid only when the concentration of RNA-containing droplets is low and fusion between such RNA-containing droplets is negligible, consistent with our observed RNA concentration of around 10^−1^ nM (Fig. 2), corresponding to ∼0.1 molecule/droplet.

### Competition assay conducted in Fig. 5b

Representative RNAs (the most frequent genotype) at each time point were used in the co-replication assay. The RNA clones were prepared as follows: The PCR amplified cDNA was ligated to a vector, pUC19 or pBAD33, using In-Fusion HD Enzyme Premix (Takara), and amplified in *E. coli* DH5α. We checked the sequence using Sanger sequencing, and if there were any unintended mutations, we corrected them using whole plasmid PCR with primers containing corrected mutations, followed by ligation and transformation as described above. The plasmids containing the correct RNA sequence were digested with Sma I (Takara) to determine the 3’-end of RNA, transcribed using T7 RNA polymerase (Takara), and purified using the RNeasy Mini Kit (QIAGEN). DNAs were degraded using DNase I (Takara) treatment, and the RNAs were purified using the RNeasy Mini Kit (QIAGEN). Each host and parasite RNA clones (1 nM) are mixed in the reconstituted translation system and incubated at 37 °C for 1 h. The concentrations of the host and parasite RNA were measured by RT-qPCR with specific primers. Primer sequences are shown in the Source Data file.

### Assay of the bottom concentration conducted in Fig. 5c

Each indicated host and parasite RNA clones were mixed at 10 nM each in the translation system, and the mixture (10 µL) was homogenized in the buffer-saturated oil phase to prepare water-in-oil droplets as described above in the coexistence experiment section. The serial dilution cycle was repeated 38–40 times under the condition close to the FR condition (dilution freqeuey of once /0.5 h and dilution ratio of 2-fold) as described in the same section.

### Calculation of generations

In the FR evolutionary experiment, the host RNA concentration was maintained for 5000 h while exchanging 20% of the reaction mixture per hour. Consequently, host RNAs underwent 1.25-fold replication per hour, resulting in 1.25^5000^-fold amplification at 5000 h. We defined one generation for an RNA replicator as the time required to double. Therefore, the total number of generations was calculated as log_2_(1.25^5000^) ∼ 1,600.

### Translation assay

The expression level of each replicase was examined using the Nano-Glo HiBiT Lytic Detection System (Promega). First, the HiBiT tag sequence was prepared by PCR using PrimeSTAR HS (Takara) with primers 31 and 32, and then, attach to the replicase gene by PCR with primer 33 and one of primers 34-36 according to its vector from plasmids constructed in the competition assay section. Then we fused these fragments by PCR with one of the primers 34-36 and primer 37. After in vitro transcription with T7 RNA polymerase (Takara), purified RNA (1 nM) was incubated at 37 ℃ for 1 h in the translation system. After translation, the solution was denatured by SDS treatment at 95 ℃ for 5 minutes in SDS buffer (50 mM Tris-HCl (pH 7.4), 2% SDS, 0.86 M 2-mercaptethanol, 10% glycerol) and diluted 10-fold with dilution buffer (50 mM Hepes-KOH (pH7.6), 100 mM KCl, 10 mM MgCl^2^, 30% glycerol, 7 mM 2-mercaptethanol, 1% BSA). An aliquot (1 µL) of the diluted samples was added to 20 µL of a reaction mixture containing HiBiT Lytic Substrate, LgBiT protein, and HiBiT Lytic Buffer, then vortexed for 4 sec and left at room temperature for 10 min. Luminescence was measured by the GloMax 20/20 Luminometer. Protein expression levels were determined according to a standard curve prepared using HiBiT control protein at concentrations ranging from 0.1 nM to 100 nM.

### Biochemical assay using purified replicases

The replicases encoded by the host species at nine time points were purified according to the previous study (Ichihashi et al. 2010) except that the KCl concentration in the buffer B was increased to 800 mM for replicases after 3984 h because they bound the column more strongly. SDS-PAGE analysis data are shown in Fig. S8. Each of these purified replicases (5 nM) and a template RNA (1 nM) was incubated in the translation system containing 30 µg/mL streptomycin to inhibit translation at 37 □ for 1 h, and RNA concentration was measured by RT-qPCR with the corresponding primer set of primer 25-26, 27-28, and 29-30.

### Calculation of the population size of the intact host RNA

During the long-term evolutionary experiment shown in Fig. 2, the host RNA concentration oscillated. The host RNA concentrations at the bottoms were reduced to approximately 10^−5^ nM at some time points (e.g., around 3000–3100 h and 4600–4900 h). These bottom concentrations correspond to approximately 5,000 molecules in a reaction tank. The RNA replicase used in this system is erroneous, and thus the population contains quasispecies that have lost replication activities. According to our previous study (see Fig. 4, Rep-RNA in (Mizuuchi and Ichihashi 2018), the frequency of host RNAs that retain more than 80% of replication activity is less than 10%. In addition, the host RNA measured here included partially degraded RNA because the quantification method used here (RT-qPCR) detects only a part (∼10%) of the host sequence. Considering that host RNA degrades at the rate of 0.1/h in the translation system (Fig. S10) and the host RNA stopped replication for approximately 40 h before reaching the bottoms of oscillations (Fig. 2), approximately 2% of host RNA, corresponding to ∼10 host RNA molecules, would be intact at the bottoms. This rough estimate supports the possibility of the occurrence of both stochastic extinction and genetic drift.

## Supplementary Materials

Supplemental Information and Source Data for this article are available.

## Acknowledgements

We thank Mrs. Kayo Aoyama and Ayu Saito for technical support, and Dr. Yuki Kanai for useful discussion.

## Author contributions

TY performed the evolutionary experiments, and KY designed and performed the other experiments. RM and NI design experiments. NI and KY wrote the manuscript.

## Data and materials availability

The data are available upon request from the authors.

