## Supplemental information for "Experimental evolution toward extinction in a molecular host-parasite system"

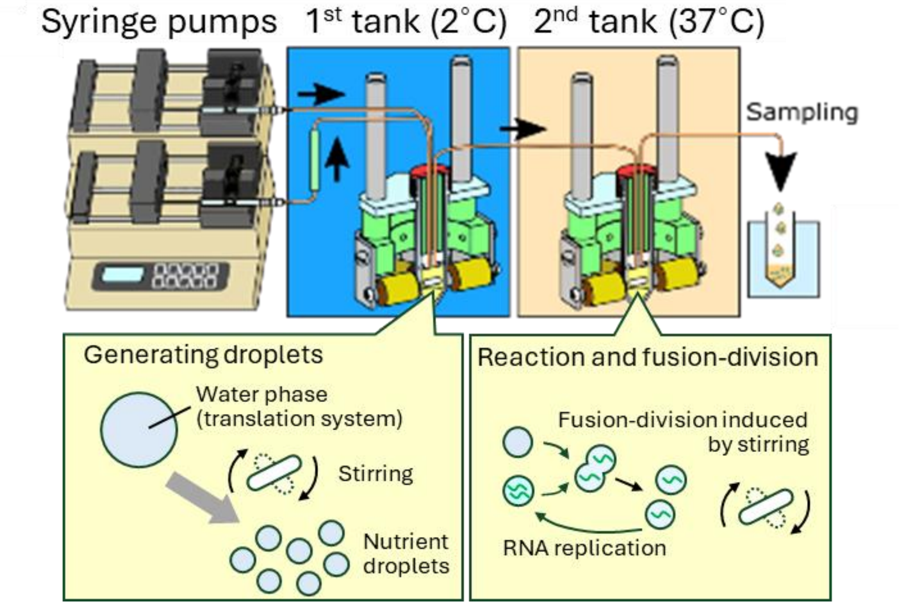


**Figure S1: Droplet flow reactor system used in this study.**

The droplet flow reactor system consisted of two 505 μL reaction tanks, each containing a magnetic stirrer. In the first tank, the oil phase and the water phase containing the reconstituted translation system were supplied separately by syringe pumps, and droplets were produced by stirring using the magnetic stirrer. The droplets were then transferred to the second reaction tank, where some droplets contained the host and parasitic RNAs, incubated at 37°C, and their replication occurred there. In the second tank, the droplets were mixed for 3 sec at 4800 rpm every 60 sec using the stirrer bar to induce fusion and division of the droplets. The droplets in the second tank were exchanged at 20% of the tank volume per hour. The illustration was modified from the previous one (Yoshiyama et al. 2016).

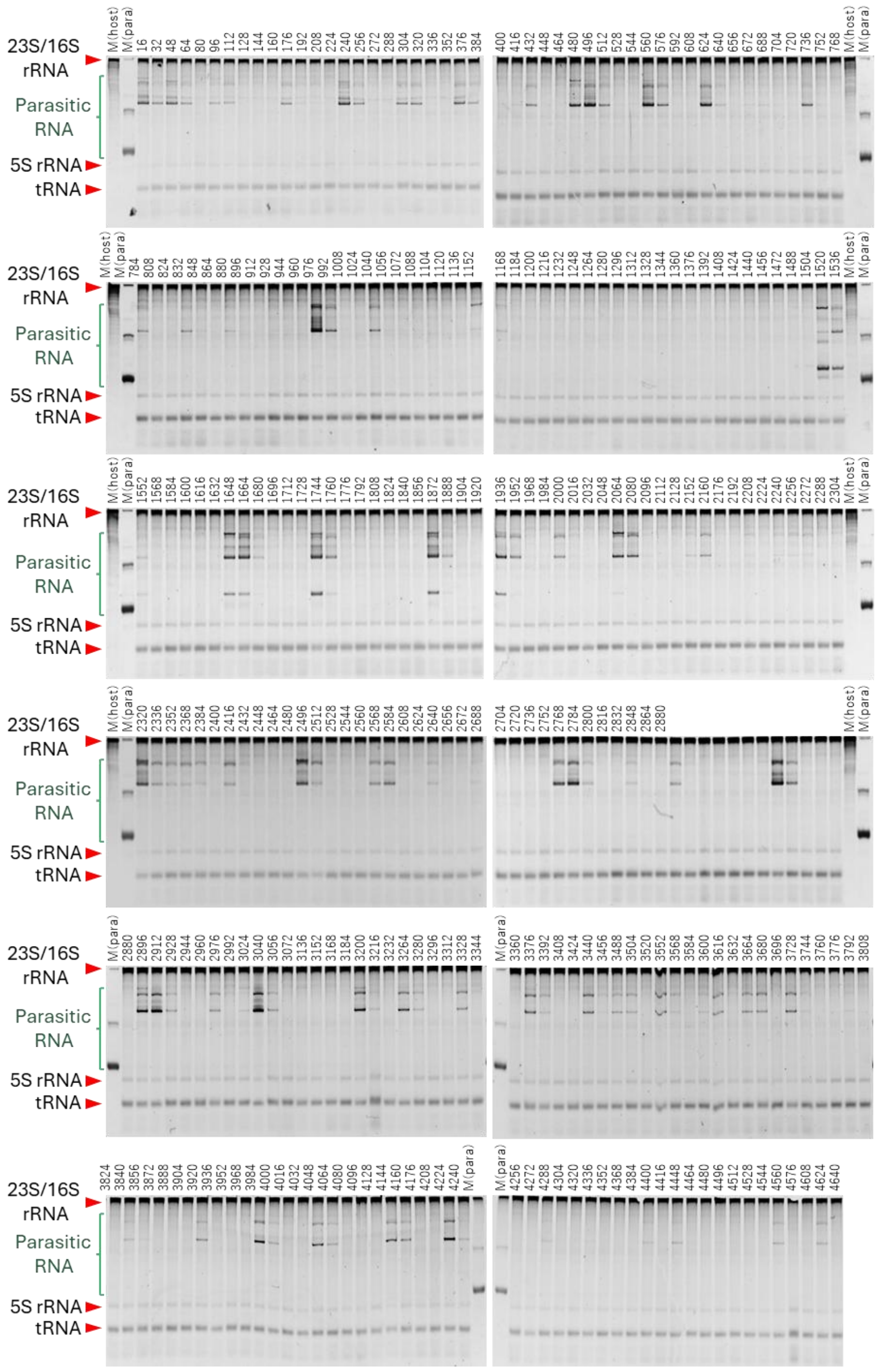


**Figure S2: Measurement of parasitic RNA concentrations.**

Purified RNA during the FR evolutionary experiment at each time point was subjected to polyacrylamide-gel electrophoresis with two size markers, a host RNA (M(host)) and a short parasite

RNA (M(para)), followed by staining with SYBR green II. All bands that appeared between 23S/16S rRNA and 5S rRNA were quantified as parasitic RNA.

| ID | Mutation | AA change | H16h | H480h | H1152h | H1520h | H2064h | H2848h | H3376h | H3984h | H4616h | New |
|---|---|---|---|---|---|---|---|---|---|---|---|---|
| 1 | G35T | | 0.00 | 0.00 | 0.00 | 0.00 | 0.00 | 0.00 | 0.00 | 0.00 | 0.36 | ○ |
| 4 | A49G | | 0.00 | 0.95 | 0.99 | 0.97 | 0.99 | 1.00 | 1.00 | 1.00 | 1.00 | |
| 5 | A53G | | 0.00 | 0.96 | 0.99 | 0.97 | 0.98 | 1.00 | 0.99 | 0.99 | 0.45 | |
| 6 | C72T | | 1.00 | 0.04 | 0.01 | 0.02 | 0.01 | 0.00 | 0.00 | 0.00 | 0.00 | |
| 8 | T78A | | 0.00 | 0.97 | 0.99 | 0.98 | 0.99 | 1.00 | 1.00 | 1.00 | 1.00 | |
| 9 | C86T | | 1.00 | 1.00 | 1.00 | 1.00 | 0.99 | 1.00 | 1.00 | 1.00 | 1.00 | |
| 13 | T114C | | 1.00 | 1.00 | 0.98 | 1.00 | 1.00 | 1.00 | 1.00 | 1.00 | 1.00 | |
| 14 | A116G | | 0.00 | 1.00 | 1.00 | 1.00 | 1.00 | 1.00 | 1.00 | 1.00 | 1.00 | |
| 15 | -129A | | 1.00 | 0.02 | 0.00 | 0.01 | 0.01 | 0.00 | 0.00 | 0.00 | 0.00 | |
| 16 | G141A | | 0.99 | 0.01 | 0.00 | 0.01 | 0.00 | 0.00 | 0.00 | 0.00 | 0.00 | |
| 19 | C160T | | 0.00 | 0.99 | 1.00 | 1.00 | 1.00 | 1.00 | 1.00 | 1.00 | 1.00 | |
| 22 | T182C | | 0.00 | 0.00 | 0.00 | 0.00 | 0.00 | 1.00 | 1.00 | 1.00 | 1.00 | ○ |
| 23 | T200C | | 0.00 | 0.00 | 0.00 | 0.00 | 0.00 | 0.00 | 0.00 | 1.00 | 1.00 | ○ |
| 24 | A215G | | 0.00 | 0.00 | 0.00 | 0.00 | 0.00 | 0.01 | 0.00 | 0.00 | 1.00 | |
| 25 | T224C | | 0.00 | 0.00 | 0.00 | 0.00 | 0.00 | 0.00 | 0.00 | 0.00 | 1.00 | ○ |
| 26 | A225G | | 1.00 | 1.00 | 1.00 | 1.00 | 1.00 | 1.00 | 1.00 | 1.00 | 1.00 | |
| 27 | A227G | | 0.01 | 0.75 | 1.00 | 1.00 | 1.00 | 1.00 | 1.00 | 1.00 | 1.00 | ○ |
| 28 | C232T | P2S | 0.99 | 0.00 | 0.00 | 0.00 | 0.00 | 0.00 | 0.00 | 0.00 | 0.00 | |
| 29 | A243G | A5syn | 0.00 | 0.00 | 0.00 | 0.00 | 1.00 | 1.00 | 1.00 | 1.00 | 1.00 | ○ |
| 30 | C255G | N9K | 0.00 | 0.00 | 1.00 | 1.00 | 1.00 | 1.00 | 1.00 | 1.00 | 1.00 | ○ |
| 31 | C259T | L11F | 1.00 | 0.00 | 0.00 | 0.00 | 0.00 | 0.00 | 0.00 | 0.00 | 0.00 | |
| 32 | A269G | Q14R | 1.00 | 0.00 | 0.00 | 0.00 | 0.00 | 0.00 | 0.00 | 0.00 | 0.00 | |
| 33 | T271C | L15syn | 0.00 | 0.00 | 0.00 | 0.00 | 0.00 | 0.00 | 0.50 | 1.00 | 1.00 | ○ |
| 34 | A291G | T21syn | 0.00 | 0.00 | 0.00 | 0.00 | 0.00 | 1.00 | 1.00 | 1.00 | 1.00 | ○ |
| 35 | T298C | I23syn | 0.00 | 0.00 | 0.00 | 0.00 | 0.74 | 0.00 | 0.00 | 0.00 | 0.00 | ○ |
| 36 | T328C | I33syn | 0.03 | 0.00 | 0.00 | 0.00 | 0.00 | 0.00 | 0.00 | 0.00 | 1.00 | ○ |
| 37 | T359G | S44A | 0.00 | 0.84 | 1.00 | 1.00 | 1.00 | 1.00 | 1.00 | 1.00 | 1.00 | ○ |
| 38 | A403G | R58syn | 0.00 | 0.00 | 0.00 | 0.00 | 0.00 | 1.00 | 1.00 | 1.00 | 1.00 | ○ |
| 39 | T501C | V91A | 0.21 | 0.02 | 0.00 | 0.00 | 0.00 | 0.00 | 0.00 | 0.00 | 0.00 | |
| 40 | G511A | E94syn | 0.00 | 0.00 | 0.00 | 0.00 | 0.00 | 0.00 | 0.00 | 1.00 | 1.00 | ○ |
| 45 | G566A | R112syn | 1.00 | 0.00 | 0.00 | 0.00 | 0.00 | 0.00 | 0.00 | 0.00 | 0.00 | |
| 51 | -614G | | 0.12 | 0.00 | 0.00 | 0.00 | 0.00 | 0.00 | 0.00 | 0.00 | 0.00 | ○ |
| 52 | A615G | I128M | 0.52 | 0.00 | 0.00 | 0.00 | 0.00 | 0.00 | 0.00 | 0.00 | 0.00 | |
| 53 | C616G | H129D | 0.49 | 0.00 | 0.00 | 0.00 | 0.00 | 0.00 | 0.00 | 0.00 | 0.00 | |
| 55 | A628C | R133syn | 1.00 | 0.00 | 0.00 | 0.00 | 0.00 | 0.00 | 0.00 | 0.00 | 0.00 | |
| 58 | T657C | V142A | 0.00 | 0.00 | 1.00 | 0.00 | 0.00 | 0.00 | 0.00 | 0.00 | 0.00 | ○ |
| 69 | A755G | A174syn | 0.01 | 1.00 | 0.99 | 1.00 | 1.00 | 1.00 | 1.00 | 1.00 | 1.00 | |
| 70 | C756T | L175F | 0.01 | 1.00 | 1.00 | 1.00 | 1.00 | 1.00 | 1.00 | 1.00 | 1.00 | |
| 72 | T794C | V187syn | 0.00 | 0.00 | 1.00 | 1.00 | 1.00 | 1.00 | 1.00 | 1.00 | 1.00 | ○ |
| 73 | T818C | H195syn | 0.00 | 0.00 | 0.00 | 0.00 | 0.00 | 0.00 | 0.71 | 1.00 | 1.00 | ○ |
| 74 | A855G | K208E | 0.00 | 1.00 | 1.00 | 1.00 | 1.00 | 1.00 | 1.00 | 1.00 | 1.00 | |
| 75 | A857G | E208syn | 0.00 | 0.00 | 0.00 | 0.00 | 0.00 | 0.00 | 0.00 | 0.00 | 1.00 | ○ |
| 76 | -860A | | 0.10 | 0.00 | 0.00 | 0.00 | 0.00 | 0.00 | 0.00 | 0.00 | 0.00 | ○ |
| 77 | A861G | A209syn | 0.56 | 0.00 | 0.00 | 0.00 | 0.00 | 0.00 | 0.00 | 0.00 | 0.00 | |
| 78 | T891C | D219syn | 0.00 | 0.00 | 0.00 | 0.00 | 0.00 | 0.00 | 0.00 | 0.00 | 1.00 | ○ |
| 79 | A909G | E225syn | 0.00 | 0.00 | 0.00 | 0.00 | 0.00 | 0.00 | 0.93 | 1.00 | 1.00 | ○ |

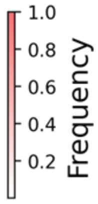


**Figure S3: Dominant host mutations and their frequencies.**

Host mutations were found at a frequency of 10% or higher at least one analyzed time point. “New” column indicates mutations not observed in the previous SD evolutionary experiment (Mizuuchi et al. 2022).

| ID | Mutation | AA change | H16h | H480h | H1152h | H1520h | H2064h | H2848h | H3376h | H3984h | H4616h | New |
|---|---|---|---|---|---|---|---|---|---|---|---|---|
| 80 | -925T | | 0.10 | 0.10 | 0.09 | 0.11 | 0.10 | 0.09 | 0.11 | 0.11 | 0.11 | ○ |
| 81 | T971C | C246R | 0.21 | 0.00 | 0.00 | 0.00 | 0.00 | 0.00 | 0.00 | 0.00 | 1.00 | |
| 82 | A1045G | L270syn | 0.00 | 0.00 | 0.00 | 0.00 | 0.00 | 0.14 | 0.00 | 0.00 | 0.00 | ○ |
| 83 | A1079G | M282V | 0.00 | 0.00 | 1.00 | 0.00 | 0.00 | 0.00 | 0.00 | 0.00 | 0.00 | ○ |
| 84 | T1096C | C287syn | 0.41 | 0.00 | 0.00 | 0.00 | 0.00 | 0.00 | 0.00 | 0.00 | 0.01 | |
| 85 | T1198C | S321syn | 0.00 | 0.00 | 0.00 | 0.00 | 0.00 | 0.00 | 0.00 | 1.00 | 1.00 | ○ |
| 86 | T1240C | I335syn | 0.00 | 1.00 | 1.00 | 1.00 | 1.00 | 1.00 | 1.00 | 1.00 | 1.00 | ○ |
| 87 | T1286C | S351P | 0.00 | 1.00 | 1.00 | 1.00 | 1.00 | 1.00 | 1.00 | 1.00 | 1.00 | |
| 88 | C1297T | V354syn | 0.00 | 0.48 | 1.00 | 1.00 | 1.00 | 1.00 | 1.00 | 1.00 | 1.00 | ○ |
| 89 | T1357C | V374syn | 0.01 | 1.00 | 1.00 | 1.00 | 1.00 | 1.00 | 1.00 | 1.00 | 1.00 | ○ |
| 90 | T1366C | Y377syn | 0.00 | 0.00 | 1.00 | 1.00 | 1.00 | 1.00 | 1.00 | 1.00 | 1.00 | ○ |
| 91 | T1375C | F380syn | 0.00 | 0.00 | 0.00 | 0.00 | 0.00 | 0.00 | 1.00 | 1.00 | 1.00 | ○ |
| 92 | T1387C | T384syn | 0.00 | 0.00 | 0.00 | 0.00 | 0.00 | 0.15 | 0.00 | 0.00 | 0.00 | ○ |
| 93 | T1485C | V417A | 0.00 | 0.00 | 1.00 | 1.00 | 1.00 | 1.00 | 1.00 | 1.00 | 1.00 | ○ |
| 94 | G1486A | A417syn | 0.00 | 0.00 | 0.00 | 0.00 | 0.00 | 0.00 | 0.00 | 0.00 | 1.00 | ○ |
| 95 | T1492C | P419syn | 0.00 | 1.00 | 1.00 | 1.00 | 1.00 | 0.99 | 1.00 | 1.00 | 1.00 | ○ |
| 96 | T1574C | Y447H | 1.00 | 0.00 | 0.00 | 0.00 | 0.00 | 0.00 | 0.00 | 0.00 | 0.00 | |
| 97 | T1578G | L448R | 0.00 | 1.00 | 1.00 | 1.00 | 1.00 | 1.00 | 1.00 | 1.00 | 1.00 | |
| 98 | A1601G | K456E | 0.00 | 1.00 | 1.00 | 0.99 | 1.00 | 1.00 | 1.00 | 1.00 | 1.00 | |
| 99 | A1605G | Q457R | 0.00 | 1.00 | 1.00 | 1.00 | 1.00 | 1.00 | 1.00 | 1.00 | 1.00 | ○ |
| 100 | A1611G | Q459R | 0.00 | 1.00 | 1.00 | 1.00 | 1.00 | 1.00 | 1.00 | 1.00 | 1.00 | |
| 101 | A1614G | H460R | 0.01 | 1.00 | 1.00 | 1.00 | 1.00 | 1.00 | 1.00 | 1.00 | 1.00 | |
| 102 | T1639C | G468syn | 0.00 | 0.02 | 0.00 | 0.00 | 0.00 | 0.01 | 0.00 | 1.00 | 1.00 | |
| 103 | T1668C | I478T | 0.35 | 0.00 | 0.00 | 0.00 | 0.00 | 0.00 | 0.00 | 0.00 | 0.00 | |
| 104 | -1682A | | 0.00 | 0.13 | 0.13 | 0.14 | 0.13 | 0.13 | 0.13 | 0.15 | 0.10 | ○ |
| 105 | A1683G | K483E | 0.95 | 0.00 | 0.00 | 0.00 | 0.00 | 0.00 | 0.00 | 0.00 | 0.00 | |
| 106 | G1694A | G486syn | 0.00 | 0.83 | 0.00 | 1.00 | 1.00 | 1.00 | 1.00 | 1.00 | 0.99 | ○ |
| 107 | T1695G | W487G | 0.00 | 1.00 | 1.00 | 1.00 | 1.00 | 0.99 | 1.00 | 1.00 | 1.00 | ○ |
| 108 | G1697A | G487syn | 0.00 | 0.00 | 0.00 | 0.00 | 0.00 | 0.00 | 0.00 | 1.00 | 1.00 | ○ |
| 109 | G1732A | R499K | 0.00 | 0.00 | 0.00 | 0.00 | 0.00 | 1.00 | 1.00 | 0.99 | 1.00 | ○ |
| 110 | A1738G | Q501R | 0.00 | 0.00 | 0.00 | 0.00 | 0.00 | 1.00 | 0.98 | 1.00 | 1.00 | ○ |
| 111 | C1745T | R503syn | 0.00 | 1.00 | 1.00 | 0.99 | 1.00 | 0.99 | 0.99 | 0.00 | 0.00 | |
| 112 | T1785C | C517R | 0.00 | 0.00 | 0.00 | 0.00 | 0.00 | 1.00 | 1.00 | 1.00 | 1.00 | ○ |
| 113 | G1819T | R528M | 0.00 | 1.00 | 1.00 | 1.00 | 1.00 | 0.99 | 0.99 | 0.99 | 0.98 | |
| 114 | T1833C | C533R | 0.00 | 0.00 | 0.00 | 0.00 | 0.00 | 1.00 | 1.00 | 1.00 | 0.98 | ○ |
| 115 | T1839C | F535L | 0.00 | 0.00 | 0.00 | 0.00 | 0.00 | 0.00 | 0.00 | 1.00 | 0.98 | ○ |
| 116 | T1847C | D537syn | 0.00 | 0.00 | 0.00 | 0.00 | 0.00 | 0.00 | 0.00 | 1.00 | 0.98 | ○ |
| 117 | T1910C | G558syn | 0.00 | 0.00 | 1.00 | 1.00 | 1.00 | 1.00 | 1.00 | 1.00 | 0.98 | ○ |
| 118 | T1915A | F560Y | 0.00 | 1.00 | 1.00 | 1.00 | 1.00 | 1.00 | 1.00 | 1.00 | 0.98 | ○ |
| 119 | A1918G | D561G | 0.00 | 0.00 | 0.00 | 0.00 | 0.00 | 0.00 | 0.00 | 1.00 | 0.98 | ○ |
| 120 | A1920G | I562V | 0.00 | 0.00 | 0.00 | 0.00 | 0.00 | 0.00 | 0.00 | 1.00 | 0.98 | ○ |
| 121 | T1943A | C569* | 0.00 | 0.00 | 1.00 | 1.00 | 1.00 | 1.00 | 1.00 | 1.00 | 0.98 | ○ |
| 122 | T1952C | C572syn | 0.18 | 0.00 | 0.00 | 0.00 | 0.00 | 0.00 | 0.00 | 0.00 | 0.00 | |
| 123 | T1953C | C573R | 0.33 | 0.00 | 0.00 | 0.00 | 0.00 | 0.00 | 0.00 | 0.00 | 0.00 | |
| 125 | A1980C | | 0.00 | 0.91 | 0.99 | 0.99 | 1.00 | 1.00 | 1.00 | 1.00 | 0.98 | |
| 128 | A1991T | | 0.00 | 0.98 | 0.99 | 0.99 | 1.00 | 1.00 | 1.00 | 1.00 | 0.98 | ○ |

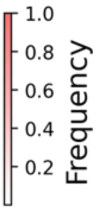


**Figure S3 (continued)**

| ID | Mutation | P16h | P480h | P1152h | P1520h | P2064h | P2848h | P3376h | P3984h | New |
|---|---|---|---|---|---|---|---|---|---|---|
| 2 | A40G | 0.00 | 0.00 | 0.00 | 0.00 | 0.07 | 0.65 | 0.84 | 0.93 | |
| 3 | C41T | 0.00 | 0.00 | 0.00 | 0.00 | 0.06 | 0.15 | 0.44 | 0.56 | ○ |
| 4 | A49G | 0.00 | 0.00 | 0.00 | 0.01 | 0.01 | 0.71 | 0.97 | 0.95 | |
| 6 | C72T | 0.99 | 1.00 | 1.00 | 0.99 | 0.99 | 0.07 | 0.02 | 0.04 | |
| 7 | G73A | 0.00 | 0.00 | 0.00 | 0.00 | 0.00 | 0.00 | 0.01 | 0.25 | ○ |
| 9 | C86T | 1.00 | 1.00 | 1.00 | 1.00 | 1.00 | 1.00 | 1.00 | 1.00 | |
| 10 | C88T | 0.13 | 1.00 | 1.00 | 0.99 | 1.00 | 0.97 | 0.98 | 0.99 | |
| 11 | A92G | 0.00 | 0.10 | 0.96 | 0.97 | 0.99 | 0.96 | 0.96 | 0.97 | ○ |
| 12 | G95A | 0.52 | 0.99 | 1.00 | 0.99 | 0.99 | 0.96 | 0.97 | 0.99 | |
| 13 | T114C | 1.00 | 0.99 | 0.99 | 1.00 | 1.00 | 1.00 | 1.00 | 0.99 | |
| 14 | A116G | 0.00 | 1.00 | 1.00 | 1.00 | 1.00 | 1.00 | 1.00 | 1.00 | |
| 15 | -129A | 1.00 | 1.00 | 1.00 | 1.00 | 1.00 | 0.99 | 0.99 | 0.99 | |
| 16 | G141A | 1.00 | 0.99 | 0.99 | 0.98 | 0.95 | 0.99 | 0.99 | 0.98 | |
| 17 | C160A | 0.00 | 0.02 | 0.12 | 0.04 | 0.01 | 0.00 | 0.01 | 0.00 | |
| 18 | C160G | 0.13 | 0.00 | 0.00 | 0.00 | 0.00 | 0.00 | 0.00 | 0.00 | ○ |
| 20 | G172A | 0.56 | 1.00 | 1.00 | 1.00 | 1.00 | 1.00 | 1.00 | 1.00 | |
| 21 | G178C | 0.99 | 1.00 | 0.99 | 1.00 | 0.99 | 1.00 | 1.00 | 1.00 | |
| 41 | T532C | 0.00 | 0.01 | 0.00 | 0.00 | 0.00 | 0.00 | 0.00 | 0.13 | ○ |
| 42 | A534G | 0.16 | 0.31 | 0.98 | 0.95 | 1.00 | 0.99 | 1.00 | 0.98 | |
| 43 | T541C | 0.01 | 0.10 | 0.01 | 0.03 | 0.00 | 0.00 | 0.00 | 0.02 | ○ |
| 44 | -564G | 0.00 | 0.02 | 0.12 | 0.03 | 0.17 | 0.01 | 0.03 | 0.02 | ○ |
| 45 | G566A | 1.00 | 0.99 | 0.87 | 0.78 | 0.86 | 0.07 | 0.03 | 0.05 | |
| 46 | T584C | 0.00 | 0.00 | 0.00 | 0.00 | 0.00 | 0.93 | 0.94 | 0.96 | ○ |
| 47 | C587A | 0.01 | 0.00 | 0.00 | 0.00 | 0.01 | 0.00 | 0.02 | 0.15 | ○ |
| 48 | C587T | 0.22 | 0.00 | 0.01 | 0.00 | 0.01 | 0.01 | 0.00 | 0.05 | |
| 49 | A588G | 0.00 | 0.00 | 0.82 | 0.99 | 0.99 | 0.98 | 0.97 | 0.93 | ○ |
| 50 | T611A | 0.97 | 1.00 | 1.00 | 1.00 | 1.00 | 1.00 | 1.00 | 1.00 | |
| 52 | A615G | 0.00 | 0.00 | 0.00 | 0.00 | 0.00 | 0.02 | 0.03 | 0.47 | |
| 54 | C616T | 0.01 | 1.00 | 1.00 | 1.00 | 1.00 | 0.99 | 1.00 | 1.00 | ○ |
| 55 | A628C | 1.00 | 1.00 | 1.00 | 1.00 | 1.00 | 1.00 | 1.00 | 1.00 | |
| 56 | C638T | 0.00 | 0.00 | 0.20 | 0.64 | 0.87 | 0.00 | 0.00 | 0.02 | ○ |
| 57 | -640C | 0.00 | 0.00 | 0.07 | 0.03 | 0.13 | 0.00 | 0.00 | 0.01 | ○ |
| 59 | G670A | 0.00 | 0.21 | 0.99 | 0.96 | 1.00 | 0.99 | 1.00 | 0.98 | |
| 60 | C680- | 0.01 | 0.00 | 0.00 | 0.00 | 0.11 | 0.00 | 0.00 | 0.00 | ○ |
| 61 | C680A | 0.00 | 0.01 | 0.19 | 0.01 | 0.08 | 0.93 | 0.97 | 0.96 | ○ |
| 62 | -685C | 0.01 | 0.02 | 0.17 | 0.01 | 0.05 | 0.00 | 0.01 | 0.00 | ○ |
| 63 | T687C | 0.02 | 0.01 | 0.16 | 0.07 | 0.02 | 0.05 | 0.01 | 0.00 | |
| 64 | T696A | 0.00 | 0.01 | 0.07 | 0.02 | 0.01 | 0.02 | 0.00 | 0.50 | ○ |
| 65 | T696C | 0.99 | 0.97 | 0.92 | 0.98 | 0.97 | 0.97 | 0.99 | 0.50 | |
| 66 | A714G | 0.00 | 0.00 | 0.00 | 0.01 | 0.00 | 0.93 | 0.97 | 0.96 | ○ |
| 67 | C723T | 0.00 | 0.01 | 0.00 | 0.03 | 0.05 | 0.93 | 0.98 | 0.97 | ○ |
| 68 | T729C | 0.00 | 0.00 | 0.00 | 0.00 | 0.00 | 0.01 | 0.03 | 0.13 | ○ |
| 71 | T768C | 0.38 | 0.00 | 0.00 | 0.00 | 0.01 | 0.01 | 0.00 | 0.00 | |
| 124 | C1974T | 0.15 | 1.00 | 1.00 | 1.00 | 1.00 | 1.00 | 1.00 | 1.00 | |
| 126 | T1981C | 0.06 | 0.00 | 0.01 | 0.04 | 0.11 | 0.93 | 0.97 | 0.95 | ○ |
| 127 | -1983C | 0.00 | 0.01 | 0.01 | 0.01 | 0.12 | 0.01 | 0.02 | 0.02 | ○ |
| 129 | T2011C | 0.00 | 0.01 | 0.00 | 0.00 | 0.01 | 0.13 | 0.10 | 0.52 | ○ |
| 130 | 35-73 Del | 0.00 | 0.00 | 0.00 | 0.00 | 0.00 | 0.11 | 0.01 | 0.00 | ○ |

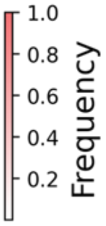


**Figure S4: Dominant parasite mutations and their frequencies**

Dominant mutations were defined as mutations that were found at a frequency of 10% or higher at least one analyzed time point. “New” column indicates mutations not observed in previous studies.

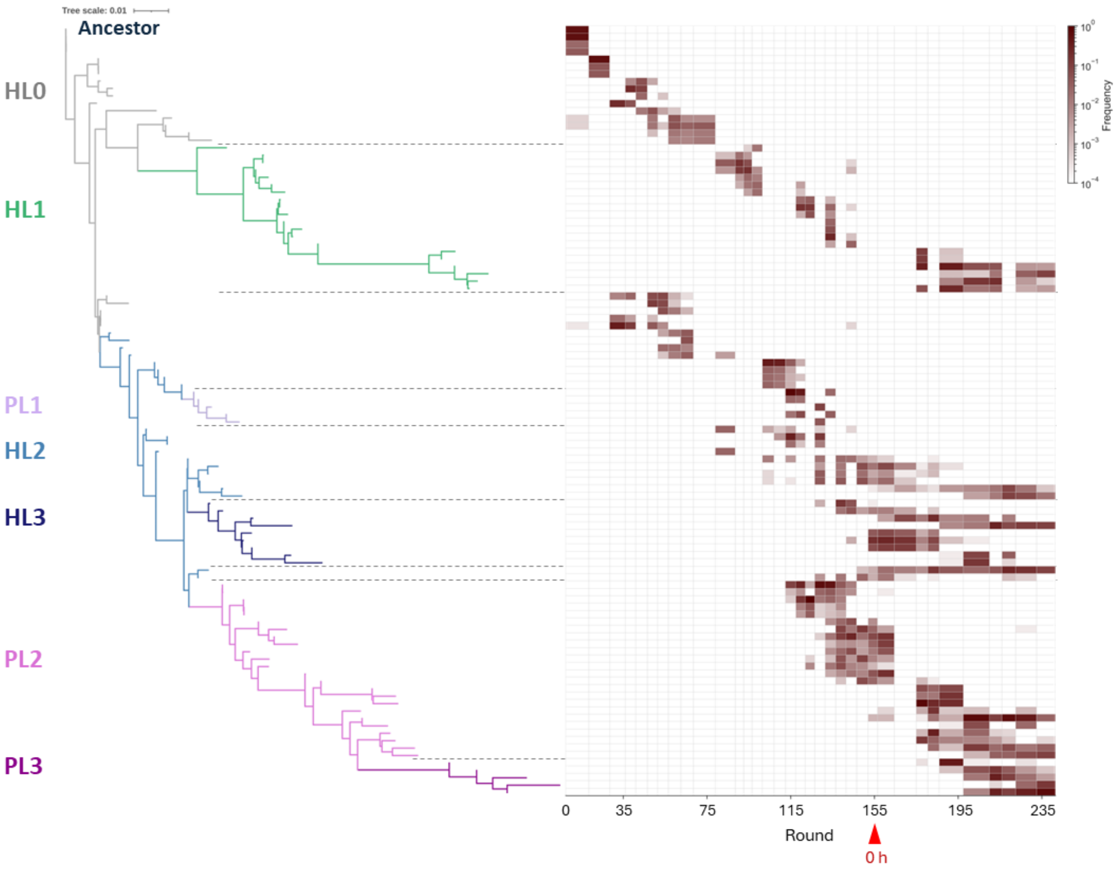


**Figure S5: Phylogenetic tree and frequencies in the population of the previous SD evolutionary experiment**

This figure was reproduced from the previous study (Mizuuchi et al. 2022). The round 155, from which the new FR evolutionary experiment started in this study, was indicated with '0 h'. In the previous SD evolutionary experiment, the five lineages existed in the final round 240.

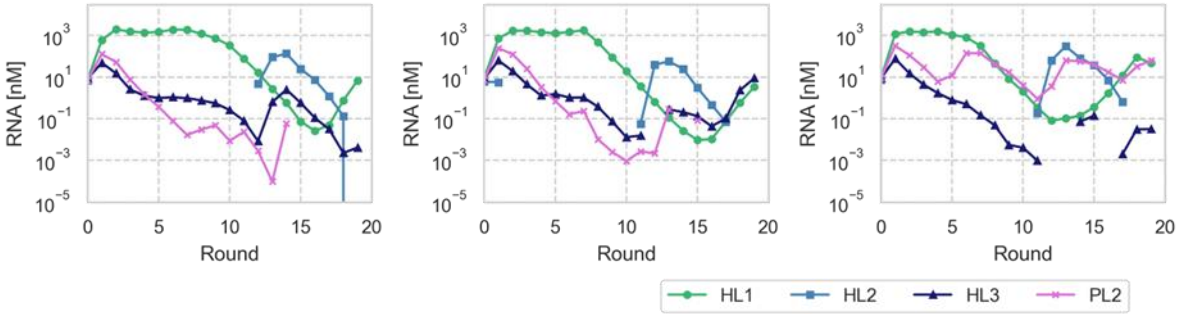


**Figure S6: Coexistence experiments using RNA species present in round 155 of the previous SD evolutionary experiment.**

Short-term serial dilution experiments were performed using three hosts (HL1, HL2, and HL3) and one parasite (PL2) species present in round 155 of the previous SD evolutionary experiment (0 h at the new FR evolutionary experiment). Four RNA clones (10 nM each) were incubated for 5 h at 37°C, then 5-fold diluted with new water-in-oil droplets using the serial dilution method. Each RNA was measured by RT-qPCR.

RNA

| Primers | HL1 | HL2 | HL3 | PL2 | PL3 |
|---|---|---|---|---|---|
| HL1 | 1 | $1.97 \times 10^{-6}$ | $1.21 \times 10^{-6}$ | $1.32 \times 10^{-7}$ | $8.61 \times 10^{-8}$ |
| HL2 | $6.68 \times 10^{-5}$ | 1 | $2.44 \times 10^{-3}$ | 0 | 0 |
| HL3 | $4.47 \times 10^{-4}$ | $6.16 \times 10^{-4}$ | 1 | 0 | $2.46 \times 10^{-7}$ |
| PL2 | $4.76 \times 10^{-5}$ | $2.80 \times 10^{-5}$ | $2.90 \times 10^{-5}$ | 1 | $1.96 \times 10^{-5}$ |
| PL3 | $8.21 \times 10^{-6}$ | $6.23 \times 10^{-8}$ | $4.26 \times 10^{-7}$ | $1.86 \times 10^{-3}$ | 1 |

RNA

| Primers | HL1-158 | H2064 | PL2-155 | P2064 |
|---|---|---|---|---|
| HL1-158 | 1 | $5.18 \times 10^{-4}$ | $9.52 \times 10^{-9}$ | 0 |
| H2064 | $8.40 \times 10^{-3}$ | 1 | $6.71 \times 10^{-8}$ | 0 |
| PL2-155 | 0 | 0 | 1 | $7.68 \times 10^{-7}$ |
| P2064 | 0 | 0 | $6.41 \times 10^{-8}$ | 1 |

RNA

| Primers | H2064 | H2848 | P2064 | P2848 |
|---|---|---|---|---|
| H2064 | 1 | $1.24 \times 10^{-5}$ | $5.44 \times 10^{-7}$ | $1.73 \times 10^{-5}$ |
| H2848 | $2.58 \times 10^{-5}$ | 1 | $3.54 \times 10^{-6}$ | $1.15 \times 10^{-4}$ |
| P2064 | 0 | 0 | 1 | $1.11 \times 10^{-5}$ |
| P2848 | 0 | 0 | $7.58 \times 10^{-6}$ | 1 |

RNA

| Primers | H2848 | H4616 | P2848 | P3984 |
|---|---|---|---|---|
| H2848 | 1 | $2.51 \times 10^{-8}$ | 0 | $1.29 \times 10^{-6}$ |
| H4616 | $5.56 \times 10^{-6}$ | 1 | 0 | 0 |
| P2848 | 0 | 0 | 1 | $1.11 \times 10^{-5}$ |
| P3984 | 0 | 0 | $1.84 \times 10^{-7}$ | 1 |

RNA

| Primers | HL1-158 | HL2-155 | HL3-155 | PL2-155 |
|---|---|---|---|---|
| HL1-158 | 1 | $2.47 \times 10^{-5}$ | $1.86 \times 10^{-5}$ | $6.36 \times 10^{-6}$ |
| HL2-155 | $7.60 \times 10^{-2}$ | 1 | $1.13 \times 10^{-6}$ | $2.08 \times 10^{-4}$ |
| HL3-155 | $1.81 \times 10^{-4}$ | $1.11 \times 10^{-2}$ | 1 | $1.94 \times 10^{-6}$ |
| PL2-155 | $1.00 \times 10^{-6}$ | $2.07 \times 10^{-4}$ | $1.80 \times 10^{-4}$ | 1 |

**Figure S7: Specificity of primers used for RT-qPCR.**

Each of five RNA species (0.1 nM based on A260) was measured using each primer set, and the mis-detected concentrations relative to the correct target RNA concentrations are shown. This result was used to correct the measured RNA concentrations in the coexistence experiments in Fig. 4b.

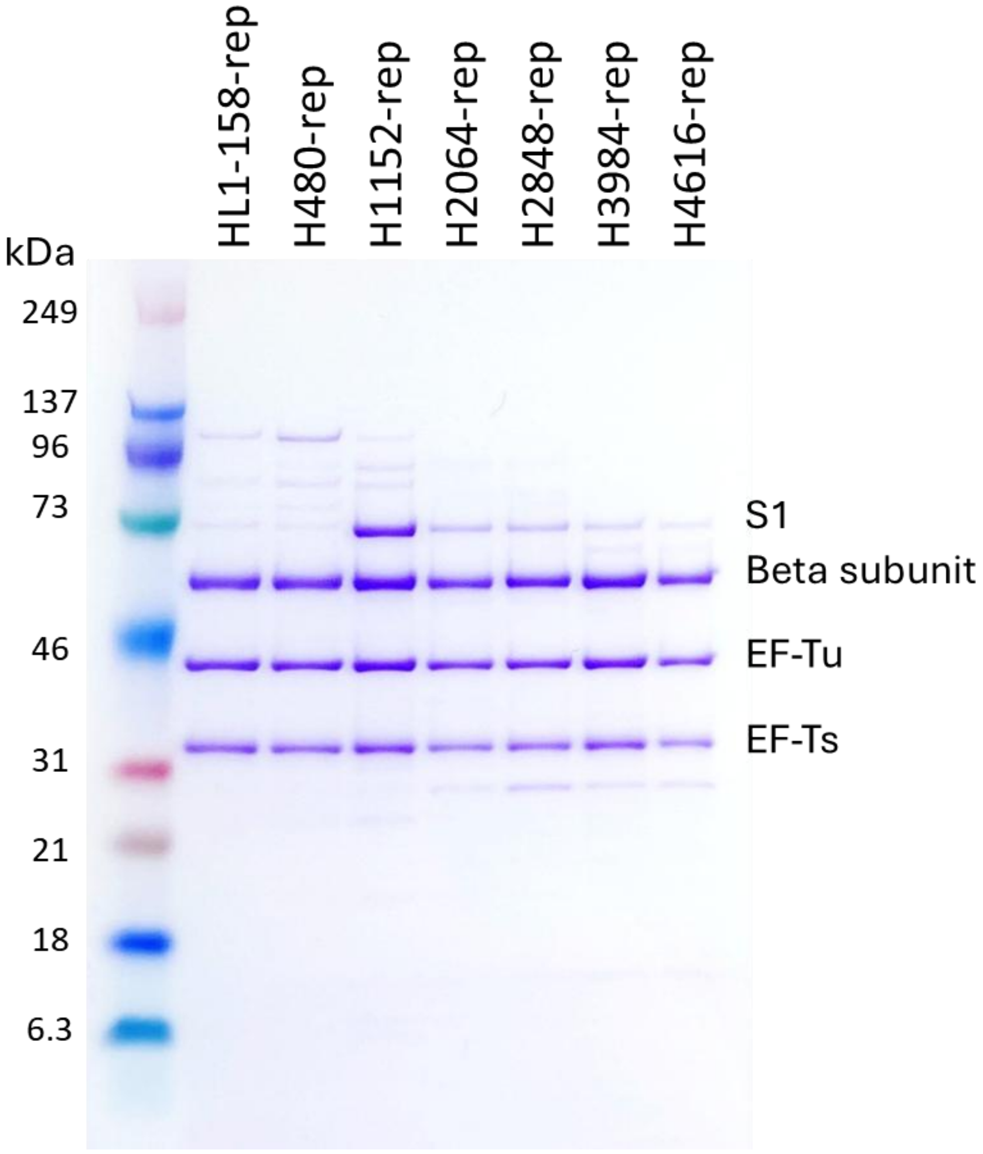


**Figure S8: SDS-PAGE analysis of recombinant replicases**

Each purified recombinant replicase (1 μg) was subjected to SDS-PAGE analysis, followed by CBB staining. S1 subunit was included in larger amounts in H1152 replicases, suggesting increased affinity for S1 for this replicase, while replication activity was not significantly different, as shown in Fig. 6.

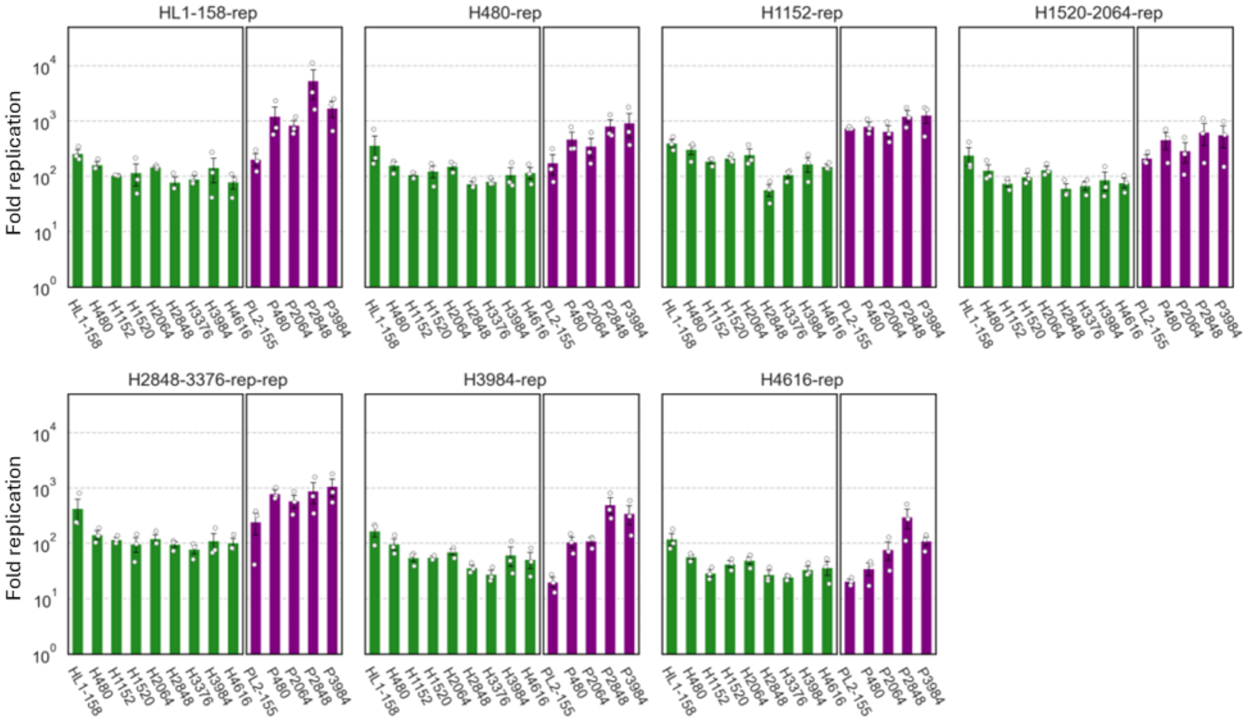


**Figure S9: Replication assay using purified replicases.**

Each purified replicase (1 nM) and each RNA species (5 nM) were incubated at 37°C for 1 h in the translation system, including streptomycin (30 μg/mL) to stop translation, and RNA concentrations were measured by RT-qPCR. The average concentrations are shown with standard errors (N = 3).

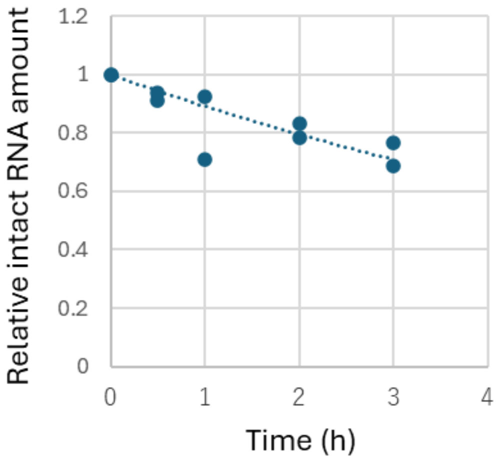


**Figure S10: RNA degradation rate in droplets.**

A $^{32}$P-labeled host RNA (70 nM), which is prepared by in vitro transcription in the presence of [α-$^{32}$P]UTP, was encapsulated in the water-in-oil droplets with a translation system without UTP to stop replication. After incubation for the indicated period at 37 ˚C, an aliquot was subjected to 8% polyacrylamide-gel electrophoresis, followed by autoradiography. The intensities of undegraded bands were measured. The regression curve (dotted line) was $y = e^{-0.115\,t}$, which means that the degradation rate was approximately 0.1 /h.

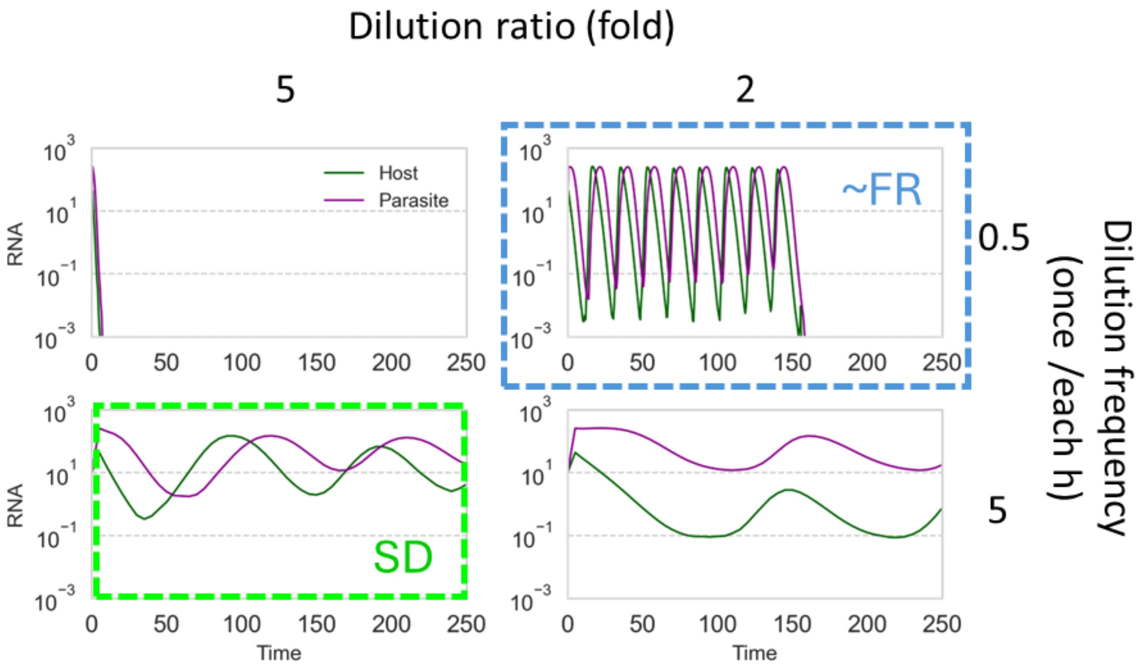


**Figure S11: Effect of dilution ratio and frequency on oscillation dynamics**.

Short-term serial dilution processes are simulated with two different dilution ratios and frequencies. Replication parameters, $k_h = 0.75$ and $k_p = 1.69$, were used. If dilution ratio was decreased from the SD condition (move rightward from the light green square), RNA concentrations at the bottoms decreased. Then, if dilution frequency was increased (move upward to light blue square), RNA concentrations at the bottoms further decreased. These results support that both small dilution ratio and higher frequency under the FR condition contribute the lower RNA concentrations at the bottoms.
